# Case Study of Single-cell Protein Activity Based Drug Prediction for Precision Treatment of Cholangiocarcinoma

**DOI:** 10.1101/2022.02.28.482410

**Authors:** Aleksandar Obradovic, Lorenzo Tomassoni, Daoqi Yu, Kristina Guillan, Katie Souto, Elise Fraser, Susan Bates, Charles G. Drake, Yvonne Saenger, Filemon Dela Cruz, Andrew Kung, Andrea Califano

## Abstract

Cholangiocarcinoma is a rare, aggressive malignancy with limited treatment options, due to a paucity of actionable mutations and low response to immune checkpoint inhibitors. Furthermore, its extreme heterogeneity prevents identification of actionable dependencies from bulk-tissue profiles. To address these challenges, we introduce a highly generalizable, single-cell framework for the mechanism-based prioritization of drugs to treat rare, highly heterogeneous tumors. Analysis of transformed cells, accounting for only 10% of a cholangiocarcinoma patient biopsy revealed three molecularly-distinct subpopulations, predicted to be sensitive to four drugs by regulatory network-based analysis. Validation in a low-passage, patient-derived xenograft (PDX) from the same patient confirmed tumor growth rate control by two of these drugs (plicamycin and dacinostat) and further validated predicted subpopulation-specific effects, suggesting they may represent promising candidates for follow-up clinical trials, either alone or in combination with current standard-of-care chemotherapies. The proposed approach can be generalized to elucidate complementary dependencies of rare, heterogeneous tumors, at the single cell level.

## Introduction

Cholangiocarcinoma (CCA) is an aggressive biliary adenocarcinoma with both intrahepatic (iCCA) and extrahepatic subtypes, the latter further characterized as either perihilar (pCCA) or distal (dCCA). Although rare, CCA accounts for up to 20% of newly diagnosed primary hepatic tumors each year, making it the second most common hepatic malignancy after hepatocellular carcinoma (HCC)^1^. It is associated with very poor prognosis, with a median survival of only 12 –37.4 months^2^. Indeed, the American Cancer Society reports that even localized disease is associated with high mortality, with 15% and 24% 5-year survival for extra- and intra-hepatic disease, respectively, and only 2% 5-year survival for metastatic disease^3^.

For patients with non-resectable disease, management differs, depending on anatomic subtype; yet no treatment option is considered curative. Localized, unresectable iCCA is sometimes treated with locoregional therapies, such as trans-arterial chemoembolization (TACE), with median overall survival of only 12-15 months. Although very few patients meet eligibility criteria, some pCCA patients may receive liver transplantation following neoadjuvant chemotherapy, increasing 5-year disease-free survival rates to as much as 65%^2^. However, the majority of patients with non-resectable or advanced disease undergoes cytotoxic chemotherapy with gemcitabine-cisplatin as first line therapy, resulting in a median survival of only 11 months, based on multiple reports^2^.

Despite rising incidence and lack of effective therapies, studies of CCA tumor micro- environment (TME) and intratumoral heterogeneity have been limited to immunohistochemical characterization of individual stromal subpopulations, especially fibroblasts, which produce an extracellular matrix so prominent that it outweighs the tumor compartment^4^. The extent of infiltration by diverse stromal cells presents a critical challenge to transcriptional profiling from bulk tumor tissue, resulting in transcriptomes that reflect the expression profile of a mixture of stromal rather than tumor cells. When combined with lack of actionable alterations—other than IDH1 in ∼13% and 1% of intra and extrahepatic cases, respectively^5^—this severely challenges the ability to use RNA-based approaches to stratify these tumors and to discover novel pharmacologic targets. Critically, while cholangiocarcinoma may best recapitulate them, these challenges are common across a variety of additional rare, aggressive, and highly heterogeneous tumors.

Single-cell RNA sequencing (scRNA-seq) has recently emerged as a valuable tool to characterize the diverse cellular subpopulations that comprise the tumor microenvironment (TME)^6, 7^. Compared to bulk-tissue RNA-seq, scRNA-seq can roughly characterize the transcriptional state of individual TME subpopulations contributing to the emergence of specific tumor phenotypes, including rare subpopulations, whose gene expression signature would be essentially undetectable from bulk-tissue samples^8^, thus helping characterize the transcriptional heterogeneity of both transformed and stromal cells. Additionally, in contrast to antibody-based approaches, scRNA-seq generates a transcriptome-wide profile of each individual cell, without requiring proteomic marker selection and antibody optimization. While single cell profiling has been used to study tumors ranging from melanoma^9, 10^, breast cancer^11^, and glioma^12^, no single-cell studies have been performed in cholangiocarcinoma. A key goal of this manuscript is thus to show that single cell analysis, even of a single sample, can provide critical mechanistic insight into the complementary dependencies of multiple transformed cell subpopulations, thus allowing identification of effective drugs for rare, highly heterogeneous tumors.

Unfortunately, due to the limited number of mRNA molecules per cell—especially from patient-derived tumor tissue—and to the physical limits of mRNA capture efficiency, the vast majority of genes in each cell (≥ 80% on average) is undetected (*gene dropout effect*) and even detected genes are represented by a small, discrete number of mRNA reads, thus challenging downstream quantitative analyses. The VIPER algorithm successfully addresses this issue by measuring the *transcriptional activity* of a protein based on the expression of its many tissue-specific transcriptional target genes (*regulon*), including both positively regulated and repressed ones^13^. Akin to a highly multiplexed (*n* = 100) gene reporter assay, this allows accurate and reproducible measurement of protein transcriptional activity, including for transcription factors (TFs) and co-factors (co-TFs), see methods. Activity of signaling proteins (SPs), and surface markers (SMs) is also effectively measured by using their transcriptional footprint, albeit with a small loss in sensitivity^14, 15^. Single cell VIPER measurements compare favorably with antibody-based technologies, yet do not require antibody availability and optimization, thus dramatically improving identification of biologically and clinically relevant subpopulations and uncharacterized mechanisms^16–19^. Critically, single cell VIPER analysis allows adapting OncoTarget and OncoTreat—two NY Dept. of Health approved, CLIA compliant algorithms, originally developed to predict drug sensitivity from bulk-tissue RNA-seq profiles—to the analysis of single-cell profiles; see^20^ for comprehensive benchmarking of these algorithms in patient derived PDX models.

By identifying aberrantly activated proteins, for which a high-affinity inhibitor is already available, OncoTarget represents a straightforward, mutation-agnostic extension of the oncogene addiction paradigm^21^, as successfully demonstrated by recent clinical^21, 22^ and preclinical trials. In contrast, OncoTreat uses the RNA-seq profiles of tumor cells perturbed with a large library of clinically relevant drugs to identify those capable of inverting the activity of the 50 most aberrantly activated and inactivated proteins, thus inducing loss of tumor state stability and tumor cell death, as originally proposed in the context of neuroendocrine tumors^23^. Indeed, we have shown that, on average, across 20 TCGA cohorts, the top 50 most aberrantly activated and inactivated proteins represent Master Regulators (MRs) responsible for canalizing the effect of a majority of functional genomic alteration in a tumor sample^24^. Extending these algorithms to the single-cell level would thus allow identification of drugs that can target specific tumor compartments, with no confounding effects from other transformed and non-transformed subpopulations in the TME.

## Results

### Conceptual Analysis Workflow

The aim of this manuscript is to propose a highly flexible and generalizable single-cell-based precision medicine pipeline for the identification of effective drugs to treat rare heterogeneous tumors, like CCA. A conceptual flowchart summarizing the analysis workflow is shown in Figure 1. Initially, fresh tissue from tumor sample biopsy of one patient was collected for dissociation and single-cell RNA Sequencing and it was also used to establish and propagate low-passage PDX models. The combination of unsupervised clustering analysis, cell type annotation algorithms and the virtual protein activity inference performed by the VIPER algorithm successfully allowed a detailed characterization of the TME. Next, tumor cells were isolated and further sub-clustered to investigate tumor heterogeneity. Once the optimal combination of gene regulatory network and cell line model was identified, distinct tumor cells subpopulations coexisting in the same sample were processed through the OncoTarget/OncoTreat precision medicine pipeline (see methods) to prioritize a list of candidate drugs to target the different tumor compartments. Then, the top 4 selected drugs were tested *in vivo* in several low-passage PDX models originally established from the patient tumor biopsy to assess tumor growth rate control and time-to-disease extension. Finally, post-treatment scRNA-Seq data was used to validate subpopulation-specific activity of the tested drugs.

**Figure 1:**
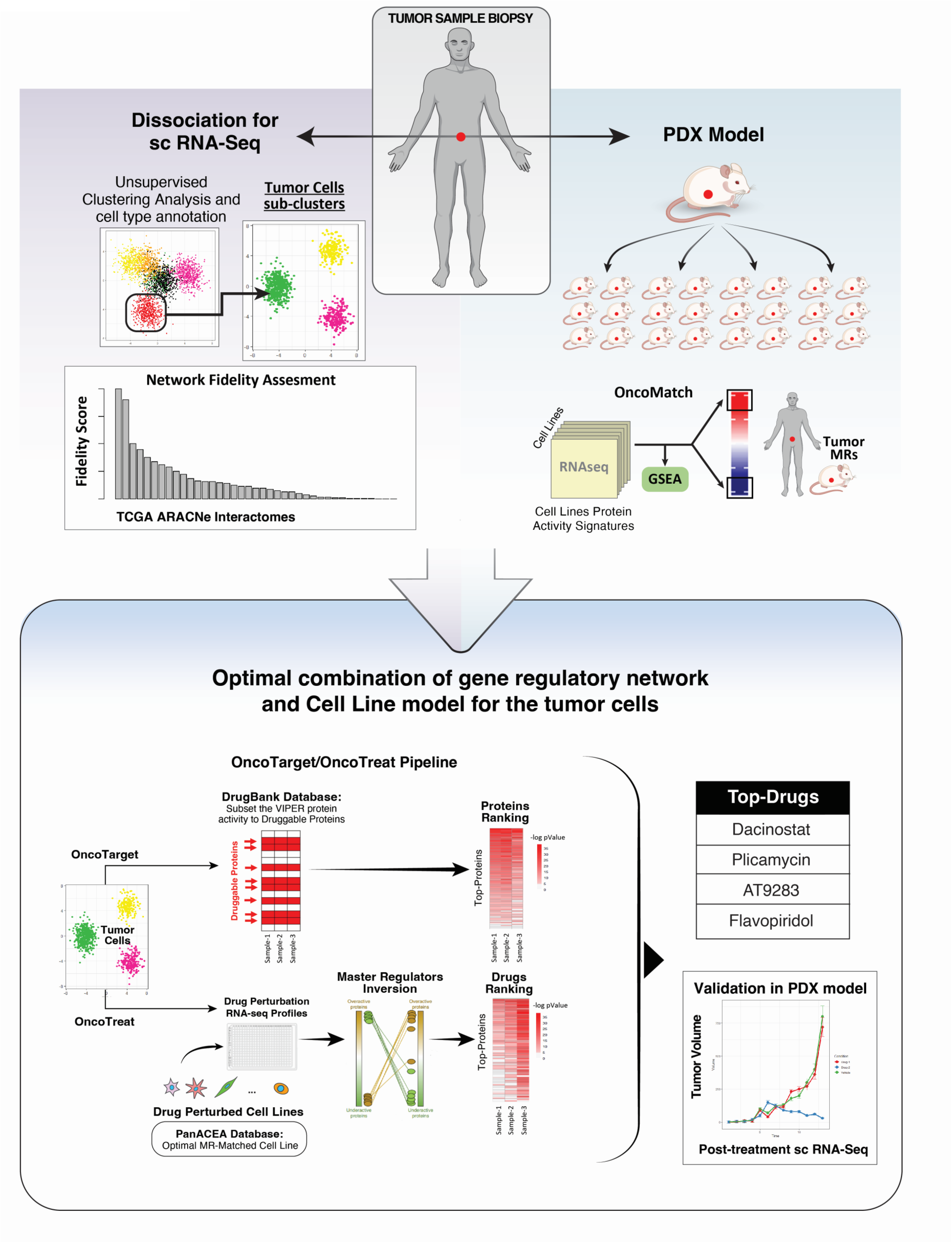
Conceptual flowchart describing an overview of the methodology proposed in the manuscript.

#### Clinical Presentation

The patient first presented to the Emergency Department with signs and symptoms of obstructive jaundice and a 6.8 cm lesion in the gall bladder fossa. Fine-needle aspiration was positive for adenocarcinoma and PET CT showed one PET avid lesion in the hilum and non-avid sub-centimeter lung nodules. Two months after initial presentation, the patient underwent a central hepatectomy with cholecystectomy and excision of bile duct tumor, which was collected for dissociation and single-cell RNA Sequencing. Pathology at that time revealed adenocarcinoma, biliary type, moderately to poorly differentiated (grades 2-3), with contiguous involvement of multiple segments of the biliary tree, including common bile duct, gall bladder, cystic duct, left and right hepatic bile ducts, liver, and perihilar soft tissue. There was also extensive lymphovascular and perineural invasion. Tumor staging was pT3N1M1, given 2/5 regional lymph nodes were positive for adenocarcinoma, and there were distant metastases noted on the falciform ligament. Immunohistochemistry was negative for HER2 overexpression, and PD-L1 combined positive score was 10. Next-generation sequencing of 467 cancer-associated genes showed a TP53 mutation as well as multiple variants of uncertain significance (EPHA5, STAT3, FAT1), with an intermediate tumor mutational burden (3.15 mutations/Mb). The tumor was microsatellite stable, and CA19-9 tumor marker levels were regularly measured to track tumor progression. The patient was initiated on combination gemcitabine, cisplatin, and paclitaxel, which were continued for seven 21-day cycles.

#### The cholangiocarcinoma TME is highly immune-infiltrated

Louvain clustering of scRNA-seq profiles revealed significant transcriptional heterogeneity, with 2,738 high-quality profiles stratified into 8 major clusters (*C*_1_ – *C*_8_) (Figure 2A-B), as defined by differentially expressed genes (Figure 2C). Cell type inference was performed by SingleR^25^, by correlating the expression of each cell with a reference database comprising RNA-seq profiles of different cellular lineages. Tumor-Infiltrating T-cells comprise the largest cluster, representing 54.3% of all cells. Based on VIPER-measured protein activity, see methods, T-cells were further stratified into (a) cytotoxic CD8 T-cells (53.5%), with high activity of cytotoxicity proteins GZMB and PRF1, as well as of the canonical marker protein CD8A; (b) CD4 T-cells (37.5%), with high CD4 and CCR7 activity, consistent with a central memory phenotype, and of VIM, a marker of tissue residency; and (c) activated CD4 T-cells (9%), presenting high STAT4 signaling activity, as well as activity of IL7R, CD3E, and IL2RB (Figure 3A-B). Notably, due gene dropouts, only CD4, VIM and IL7R could be detected as differentially expressed at the gene expression level. Protein activity-based clusters were highly concordant with SingleR-based cell type inference. Other TME subpopulations included fibroblasts (19.2%), myeloid cells (6.3%), mast cells (6.7%), endothelial cells (3.4%), B-cells (2.7%), and neutrophils (1.5%) (Figure 2, Figure 3). Consistent with this malignancy’s poor response to immune checkpoint inhibitors (https://pubmed.ncbi.nlm.nih.gov/32729929/), several of these subpopulations were previously characterized as immunosuppressive, including neutrophil myeloid-derived suppressor cells (Cluster *C*_8_)^26^ and fibroblasts (Cluster *C*_2_)^19, 27^.

**Figure 2:**
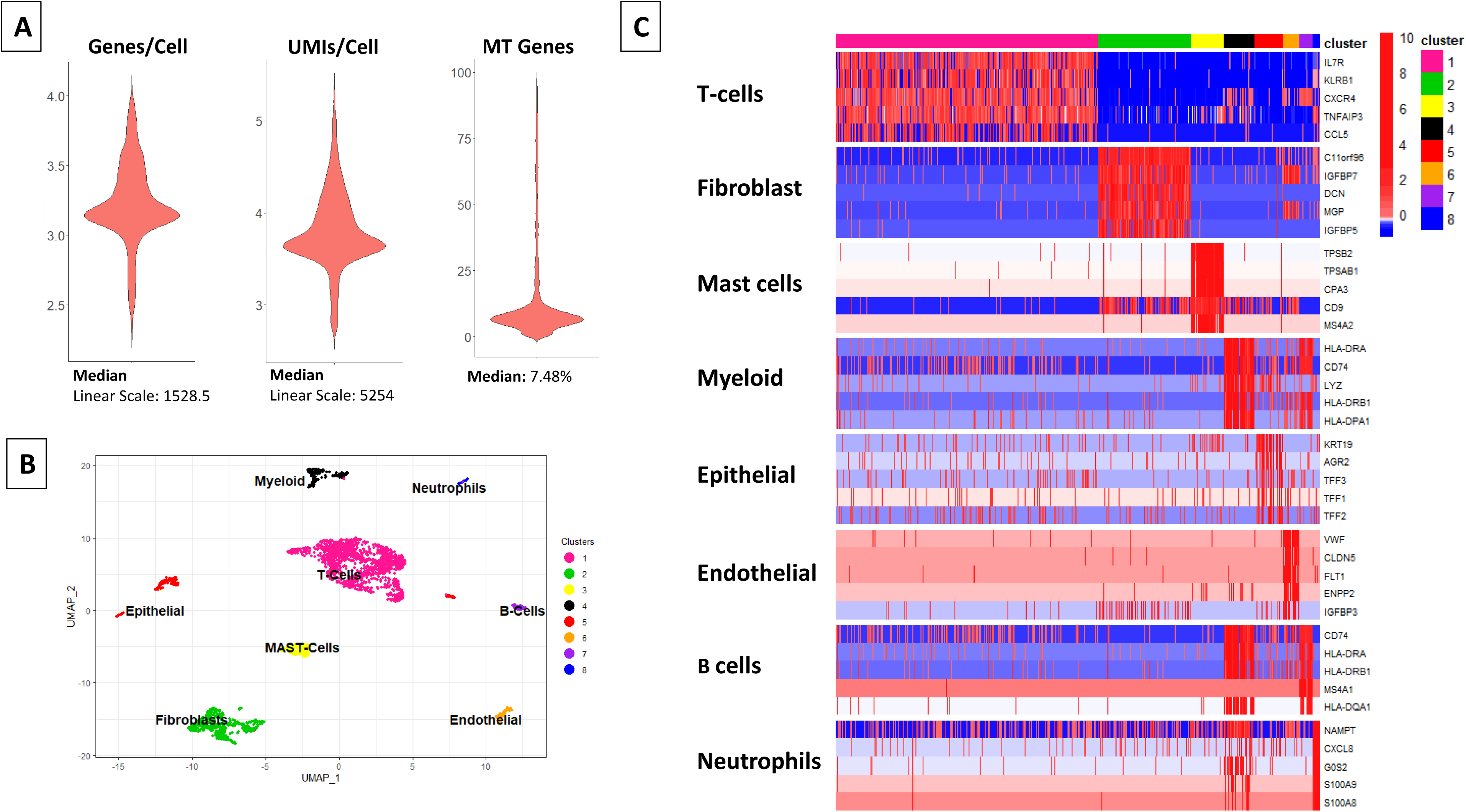
**(A)** Violin plots showing the distributions of detected genes, UMIs, and fraction of UMIs in mitochondrial (MT) genes per cell. These represent standard QC metrics used to discard low-quality cells. Y-axis of detected genes and UMIs are shown on a log-10 scale. **(B)** UMAP projection showing the results of the unsupervised, gene expression-based cluster analysis (Louvain algorithm). The cell type identity of each cluster was inferred by SingleR analysis. **(C)** Heatmap showing the top-5 most differentially upregulated genes in each cluster, as determined by comparing the average expression of each gene in one cluster versus its average expression in all other cells (MAST test)

**Figure 3:**
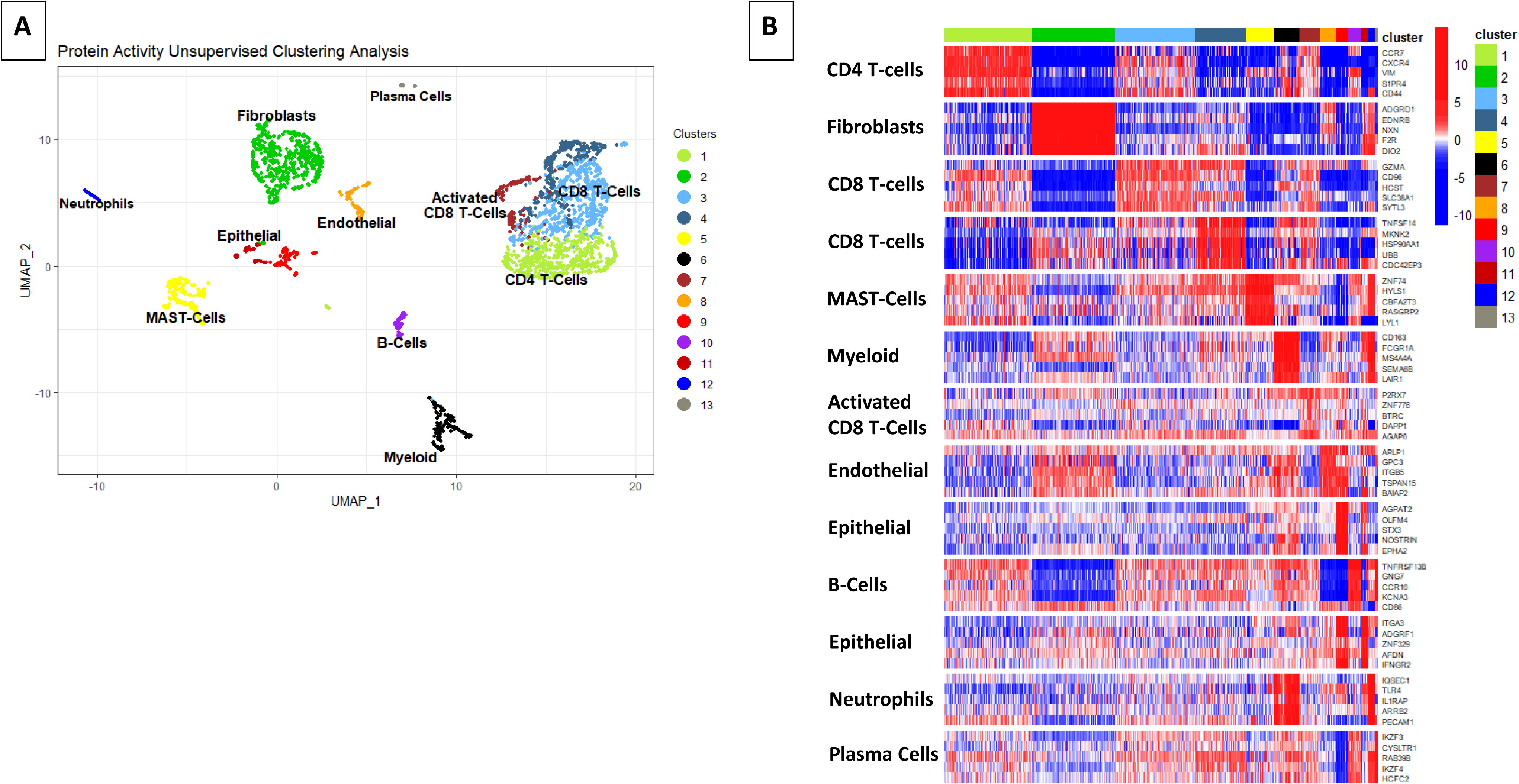
**(A)** UMAP projection of the unsupervised, protein activity-based cluster analysis, based on single-cell VIPER analysis (Louvain algorithm). **(B)** Heatmap showing the 5 most differentially active proteins in each cluster, as identified by t-Test analysis of the average activity in one cluster vs. all other clusters.

Only a relatively small population of 140 cells was identified as epithelial lineage-derived, with KRT19—an established cholangiocarcinoma marker^28^—identified as the most up-regulated gene in this population, and two canonical epithelial markers, OCLN and CDH1, aberrantly activated by VIPER analysis, yet undetected by gene expression. The transformed nature of these cells was confirmed by copy-number variation (CNV) inference (Figure 4A) (Tickle et al, 2019).

**Figure 4:**
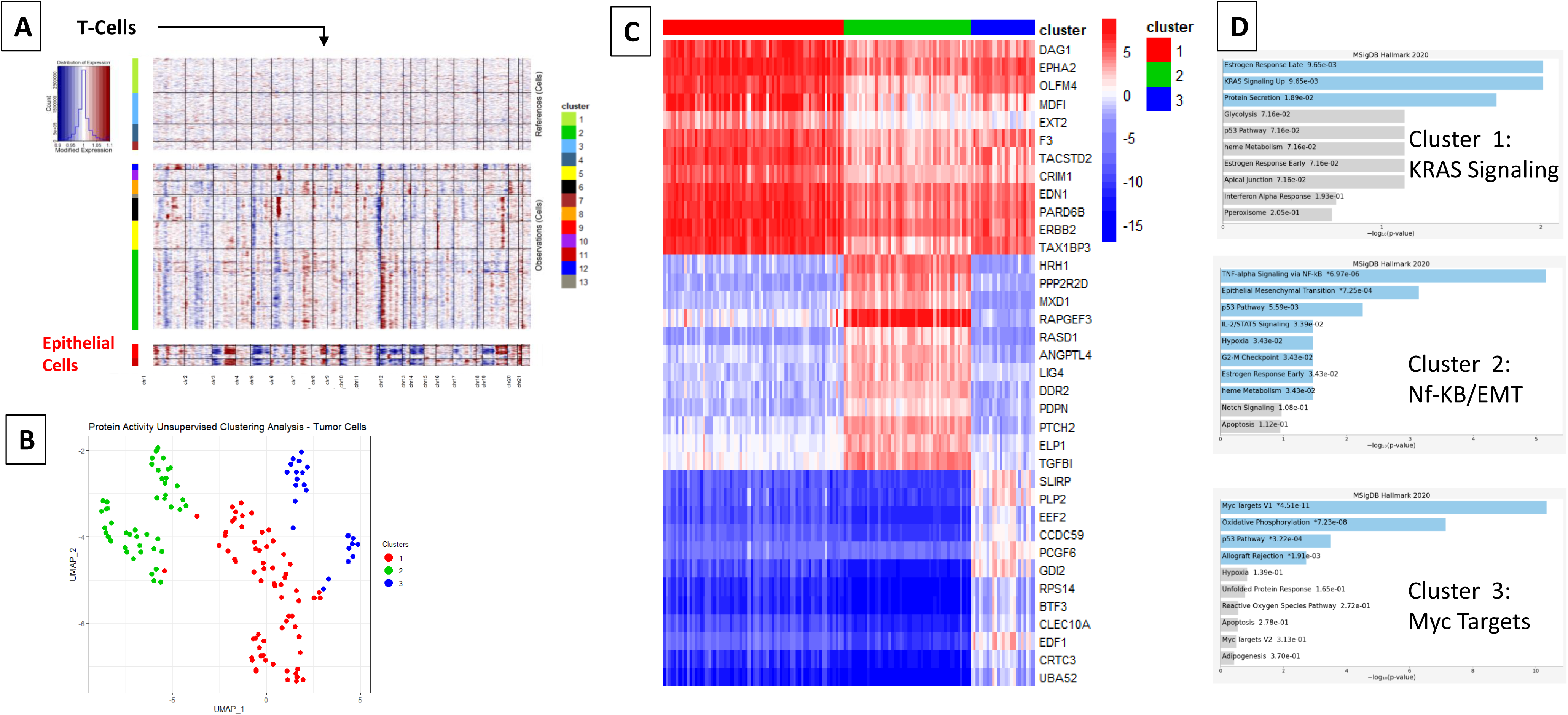
**(A)** InferCNV analysis confirmed that the epithelial cell cluster is mostly comprised of tumor cells, as they show a substantially larger complement of copy number alterations across most chromosomes, compared to T Cells chosen as a non-transformed counterpart. **(B)** UMAP projection of the 3 clusters identified from transformed cell-specific cluster analysis. **(C-D)** Heatmap showing the most differentially active proteins in each of the three tumor subpopulations identified by cluster analysis, as well as their statistically significant enriched categories from the Hallmarks of Cancer^49^.

#### Protein activity identifies molecularly-distinct, coexisting tumor cell subpopulations

Gene expression-based clustering of transformed cells failed to further stratify them, selecting a single cluster as the optimal solution. In sharp contrast, and consistent with prior studies^15, 19^, activity-based analysis—based on 1,602 transcriptional regulators, see methods—revealed three molecularly distinct subpopulations, presenting high intra-cluster homogeneity, by Silhouette analysis (Figure 4-B, Figure S4). The most differentially active proteins in each cluster are shown in Figure 4C, while the activity of key markers, including surface proteins that could be used to isolate or target cancer subsets specifically, are shown in Figure S3. Specifically (Figure 4D), the dominant cluster was characterized by aberrant activity of regulatory proteins in the KRAS signaling and estrogen response pathways, including IL6ST, FGFR3, CA12, and CDH1; a second cluster was characterized by regulatory proteins associated with upregulation of TNFa signaling via NF-kB and epithelial-mesenchymal transition (EMT), including NR4A2, NR4A3, KLF10, TNFSF9, NFKB1A, CD69, and TNF; finally, the smallest cluster was characterized by aberrant activity of proteins enriched in MYC targets and in the oxidative phosphorylation pathway, including VDAC1, ATP6V1G1, and GPI. Taken together, these data suggest that cholangiocarcinoma represents the plastic equilibrium of multiple transformed and non-transformed subpopulations.

#### Cluster-specific actionable proteins identified by single-cell OncoTarget

To perform OncoTarget and OncoTreat analysis at the single cell level, we used the average gene expression of the TCGA repository as a reference^29^, as discussed in^20^, where these methodologies were highly successful in identifying effective drugs in 34 distinct drug treatment arms in PDX models from patients who had failed multiple lines of treatment. This reference provides an effective model to assess whether a regulon is consistently differentially expressed in a specific tumor or tumor cell, compared to a large and heterogeneous collection of tumors, thus allowing identification of sample-specific, aberrantly activated and inactivated proteins.

VIPER analysis of the resulting differential gene expression signature was used to rank 5908 druggable targets, including regulatory TF/coTF and signal transduction proteins, based on their aberrant activity, see methods. Confirming the reproducibility of single cell, VIPER-based measurements, single cell OncoTarget predictions were highly consistent intra-cluster, yet diverged across clusters. Indeed, silhouette score analysis using the activity of the top statistically significant druggable proteins (p ≤ 10^-5^) confirmed prediction reproducibility within each cluster (Figure S4), suggesting that the three subpopulations identified by the analysis present distinct drug sensitivity.

Based on the reproducibility of single cell predictions, we aimed to increase sensitivity even further by adding up the mRNA read counts of all cells in a cluster (i.e., cluster-specific MetaCell) to create three cluster-specific synthetic-bulk profiles, which could then be analyzed with improved gene coverage, similar to the bulk-version of the algorithms (Figure 5E). In addition, we also generated an integrated activity signature for all proteins by integrating their VIPER-based normalized enrichment score (NES, see methods) across all cells in a cluster using Stouffer’s method (Figure 5C). Overlap of the two analyses was assessed on a cluster-by-cluster basis by measuring the correlation between the inferred protein activity of the druggable proteins resulting from the integrative analysis and from the corresponding synthetic bulk (*p*_C1_ < 2.2×10^-16^, *p*_C2_ < 2.2×10^-16^ and *p*_C3_ < 2.2×10^-16^ – Fig. 5C). . When restricted to significantly active druggable proteins, the synthetic-bulk analysis did not identify any additional targets; however, it revealed two *C*_1_ proteins (PTK2B and JAK1) as false negatives, resulting from noisy nature of scRNA-seq profiles. The almost perfect overlap of the results suggests that OncoTarget can be effectively used at the single cell level, with minimal accuracy and sensitivity losses compared to bulk profile analyses, and that potential false positives may be mitigated by generating cluster-specific synthetic bulk profiles. In addition, the analysis confirmed that, within each cluster, the activity of the proteins that control cell state (Master Regulators) is remarkably stable, from cell to cell, even though ∼80% of the genes were lost on average.

**Figure 5:**
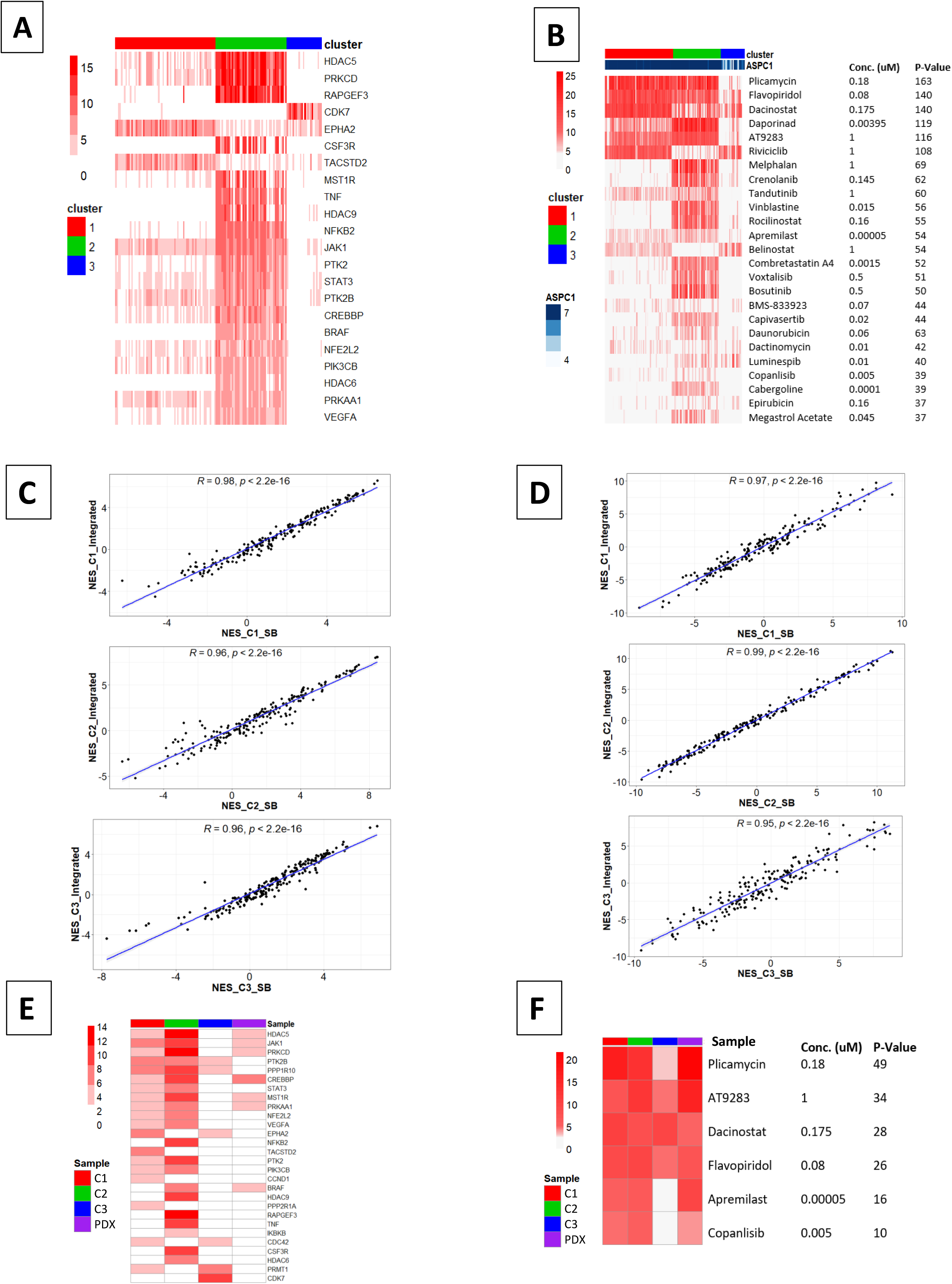
Results of OncoTarget and OncoTreat drug predictions. **(A)** OncoTarget predictions shown as the negative log_10_ *p* of the most significantly activated actionable proteins in the DrugBank database. **(B)** Negative log_10_ *p* of top OncoTreat-predicted drugs at the single cell level, tested at a concentration <= 1uM. **(C)** Comparison of OncoTarget results between the synthetic bulk and integrated analysis. X and Y axes show the VIPER NES of druggable proteins in synthetic bulk and in the integrated analysis, respectively. **(D)** Comparison of the OncoTreat results between the synthetic bulk and the integrated analysis. X and Y axes show the OncoTreat NES of each drug in the synthetic bulk and in the integrated analyses, respectively. **(E)** OncoTarget predictions from the synthetic bulks generated from each transformed subpopulation and from the PDX bulk-tissue profile. **(F)** OncoTreat predictions from the synthetic bulks generated from each transformed subpopulation and from the PDX bulk- tissue profile.

OncoTarget analysis identified 22 actionable proteins, including several histone deacetylase enzymes (HDACs), as well as other proteins of biological interest, such as the protein kinases PRKCD, PTK2, EPHA2, VEGFA, and BRAF, representing high-affinity targets of FDA-approved or late-stage (phase II/III trial) investigational compounds.

#### Cluster-specific drug sensitivity predicted by OncoTreat Analysis

We further performed drug sensitivity prediction at the single-cell and synthetic bulk level by OncoTreat, leveraging PanACEA^31^, a database of RNA-seq profiles from 16 patient-matched cell lines treated with ∼350 FDA approved and late-stage experimental (phase II/III trials) oncology drugs and vehicle control (DMSO), see methods. Cell lines were matched to patients based on Master Regulator conservation, as assessed by GSEA (OncoMatch)^23, 32^. Among the available drug perturbed cell lines, the pancreatic adenocarcinoma cell line ASPC1 presented the most significant overlap of its 50 top and bottom Master Regulator proteins (p ≤ 10^-6^, for the vast majority of transformed cells in our dataset) (Figure S1). We therefore leveraged ASPC1 as a suitable model to assess drug mechanism of action for this cholangiocarcinoma sample, as determined by the activated and inactivated proteins in drug vs. DMSO-treated cells. As described in^23^, we used OncoTreat to predict MR-inverter drugs, based on their ability to invert the activity of each single-cell’s Master Regulators. Statistical significance was assessed based on the normalized enrichment score (NES), with negative scores indicating inversion and positive scores indicating increased activity of MR proteins, as assessed by GSEA using the aREA algorithm^23^. Single-cell NES-derived p-values were integrated using Fisher’s method, such that drugs predicted to invert the MR profile of a larger number of cells, across all three clusters, were more highly ranked. The top 25 drugs emerging from this analysis are shown in Figure 5B. By performing OncoTreat analysis of cluster-specific synthetic bulk samples, drug predictions were narrowed to a set of only 6 highly significant drugs (p ≤ 10^-5^) (Figure 5F). Of these, only five (plicamycin, AT9283, dacinostat, flavopiridol, apremilast and copanlisib) were profiled at physiologically relevant concentrations (≤ 1μM).

#### OncoTreat and OncoTarget Analyses Predict Partially Overlapping Drugs

OncoTreat and OncoTarget predicted drugs significantly overlapped, with 5 of the top 25 predicted by OncoTreat also identified as high-affinity inhibitors of aberrantly activated proteins as predicted by OncoTarget (p=3.4×10^-7^).These comprise the class-I HDAC inhibitors dacinostat and belinostat—targeting HDAC5, HDAC9, and HDAC6— the HDAC6-specific inhibitor rocilinostat, the CDK7/9 inhibitor flavopiridol, and the PIK3CB inhibitor Copanlisib. Taken together, the four drugs predicted to target the greatest fraction of transformed cells across two or three clusters, at a physiologically relevant concentration (≤ 1μM), included: plicamycin, flavopiridol, dacinostat, and AT9283. Of these, dacinostat and flavopiridol were also predicted by OncoTarget analysis.

#### Bulk-tissue profile analysis fail to recapitulate single cell drug predictions

To assess the fraction of druggable target and drug candidates predicted by single-cell and cluster-based synthetic bulk analysis that would have been recapitulated by bulk profile analysis, we generated a global synthetic bulk from all single cells, by adding up their unique molecular identifier counts on a gene-by-gene basis. OncoTarget analysis failed to identify any druggable proteins (*p* ≤ 10^-5^), and only identified two shared proteins—HDAC5 (p=0.002) and CDK7 (p=0.0009)—at a much more permissive significance threshold (Figure S5A). Similarly, OncoTreat analysis on synthetic bulk of the entire sample failed to recapitulate drugs predicted to target the three tumor cell sub-clusters, with the exception of AT9283, which was the only drug concurrently predicted at single-cell level and from synthetic bulk of the entire sample with p<10^-5^(Figure S5B). These discrepancies were fully expected, because of the overwhelming majority of non-tumor cells severely diluting the molecular profile of the transformed cell subpopulations.

#### Drug validation in a Patient-Derived Xenograft (PDX) Model

We successfully engrafted and propagated a PDX model from resected tumor tissue at time of biopsy, see methods for details on IRB-approved protocol. Low-passage PDX models have high (>40%) cellularity, on average, and mouse stromal cells are eliminated from analysis as their profile does not map to the human genome. Consistently, VIPER analysis of the PDX bulk-tissue RNA-Seq presented significant enrichment of the differentially active proteins identified from the three single-cell clusters detected from single cell analysis of the patient’s sample (Figure 6A). As expected, enrichment was higher for differentially active proteins in the two larger clusters, *C*_1_ (*p* = 5.7×10^-24^) and *C*_2_ (*p* = 5.9×10^-25^), and lower for the smaller one, *C*_3_ (*p* = 5.5×10^-8^).

**Figure 6:**
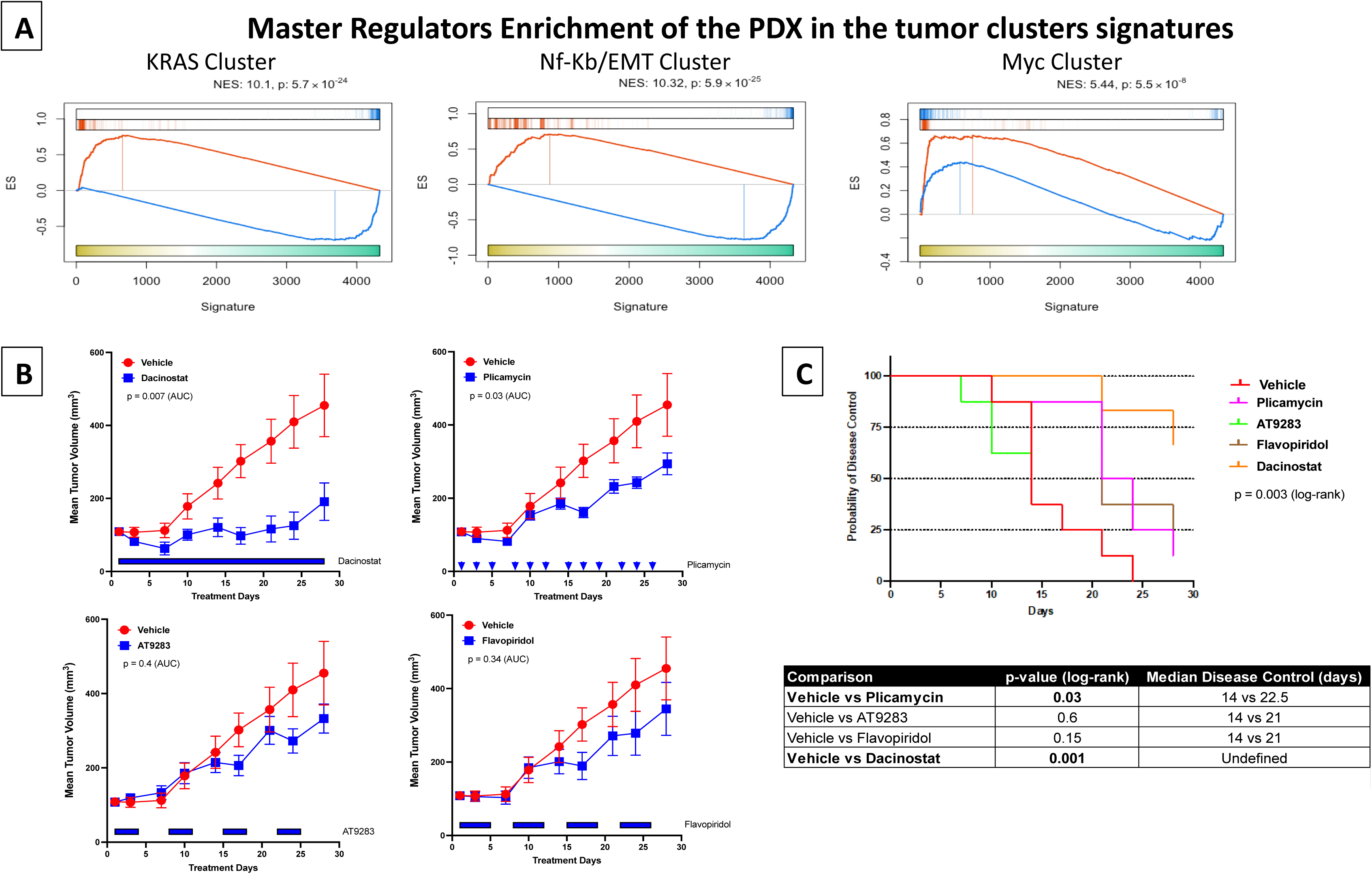
**(A)** Gene Set Enrichment Analysis (GSEA) of the top 100 and bottom 100 most differentially activated proteins (Master Regulators), for each transformed subpopulation (clusters *C*_1_ – *C*_3_), in proteins differentially active in the PDX model shows significant conservation for Clusters *C*_1_ and *C*_2_ and partial conservation (of only active Master Regulators) for *C*_3_. **(B)** Tumor growth curves comparing the volume of the tumor in PDX models treated with the 4 selected drugs vs. vehicle control. Each drug was tested in *n* = 8 mice, except for dacinostat for which we have *n* = 6. **(C)** Kaplan-Meier curve analysis assessing the time to failure (tumor volume doubling) achieved by each treatment condition.

Consistently, PDX OncoTarget analysis identified proteins that were also highly significant (*p* < 10^-5^) by OncoTarget analysis of at least one of the cluster-specific synthetic-bulk profiles (Figure 5E). Similarly, OncoTreat predicted drugs that fully overlapped between the PDX model and the two largest clusters only partially overlapped with the smallest cluster (Figure 5F). Specifically, PDX analysis identified plicamycin, AT9283, dacinostat and flavopiridol—the top four drugs with the best overall, patient-derived single cell tumor coverage (Figure 5B)—as highly significant based on PDX bulk-tissue analysis, *p*_plyc_ = 4.2×10^-22^, *p*_flavo_ = 6.6×10^-8^, *p*_AT92_ = 4.4×10^-16^, and *p*_daci_ = 1.7×10^-5^ (Figure 5F). This suggests that single cell analysis of highly heterogenous tumors can recapitulate the results from high cellularity PDX bulk tissue, while predicting subpopulation-specific effects of each drug.

The *in vivo* activity of the top four top drugs emerging from the analysis were evaluated in the patient-matched PDX model at maximum physiologically tolerated concentrations (n=8 mice/treatment arm) assessing effects of treatment on tumor growth and overall disease control. Among these drugs, dacinostat and plicamycin significantly reduced tumor growth rate (*p* = 0.007 and *p* = 0.03, respectively), with dacinostat demonstrating stable disease over 28 days of treatment (Figure 6B). Both drugs also significantly extended disease control, compared to vehicle control, based on Kaplan-Meier regression (*p*_daci_ = 0.001, with median survival time exceeding 28 days vs. 14 days, and *p*_plyc_ = 0.03, with a median survival time of 22.5 days vs. 14 days) (Figure 6C). Tumor growth analyses were performed for only 6 of 8 enrolled animals in the dacinostat treatment group due to unanticipated mortality during the study period, possibly related to drug toxicity.

Finally, to validate the predicted, subpopulation-specific activity of these drugs, we selected one animal per arm for post-treatment scRNA-seq analysis. These analyses confirmed that the molecularly distinct cellular states identified post-treatment were similar to the pre-treatment ones, suggesting that none of the drugs significantly affected the transcriptional identity of tumor cells (Figure 7A) (GSEA enrichment of pre-treatment markers in post-treatment cells: 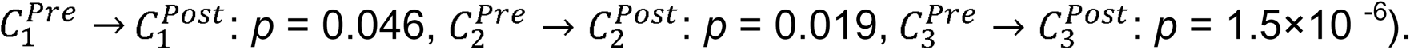.

**Figure 7:**
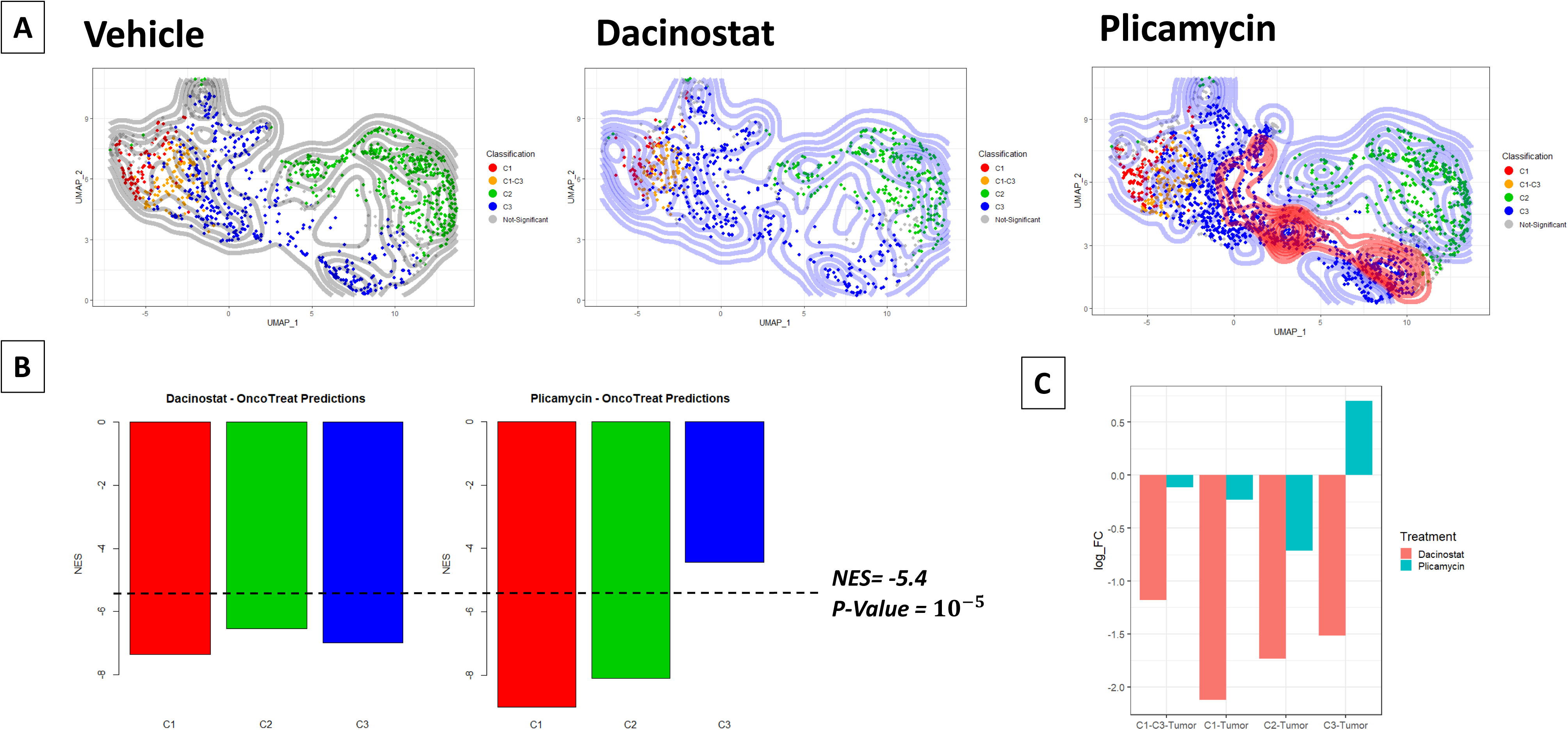
**(A)** UMAP projection of VIPER-inferred protein activity of transformed tumor cells from post-treatment PDX mice treated with dacinostat, plicamycin, and vehicle control. Differential density analysis with the respect to the vehicle is performed to assess the depletion/expansion of tumor sub-populations. Cells are colored based on the pre-treatment cluster (*C*_1_ – *C*_3_) to which they are statistically significantly matched by GSEA. **(B)** Cluster- specific, OncoTreat-predicted drug sensitivity to plicamycin and dacinostat. The y-axis shows the OncoTreat NES scores. **(C)** log_2_ Fold-Change in each transformed cell subpopulation in Figure 7B, following plicamycin and dacinostat treatment *in vivo*, relative to vehicle control, shows depletion of all subpopulations for dacinostat and depletion/increase in *C*_1_/*C*_2_ vs. *C*_3_, respectively, for plicamycin. P-values were assessed by Bonferroni corrected Fisher’s exact test with. * indicates p-value < 0.05, ** indicates p-value <0.01, *** indicates p-value <0.001

In sharp contrast, the fraction of cells in each cluster was highly affected by drug treatment, consistent with single cell OncoTreat predictions. For instance, dacinostat was predicted to invert MR protein activity for cells in all three clusters, with equivalent statistical significance (p ≤ 10^-6^). Consistently, post-treatment, PDX-based scRNA-Seq profiles showed equivalent depletion of all three transformed subpopulations, compared to vehicle control as reference (*p* < 10^-4^) (Figure 7A/C). In contrast, plicamycin was predicted to target cluster *C*_1_ and *C*_2_ cells with high specificity (*p* = 3.9×10^-17^ and *p* = 9.5×10^-14^, respectively) yet *C*_3_ cells at >10-orders of magnitude lower specificity (*p* = 1.5×10^-3^) (Figure 7B). Consistent with these predictions, post-treatment analysis of plicamycin treated PDX tumors revealed significant *C*_2_ cluster depletion and lack of *C*_1_ cluster expansion, compared to vehicle control (p_C1/Dep_ = 0.812, odds ratio *OR* = 0.96; p_c2/Dep_ = 6.03×10^-8^, *OR* = 0.643) yet significant *C*_3_ cluster expansion (p_c3/Exp_ = 3.48×10^-30^, *OR* = 2.19) (Figure 7A/C), confirming the subpopulation-specific prediction of drug sensitivity by OncoTreat analysis.

## Discussion

Reliance on statistical association methodologies requires large tumor cohorts to assess significance of recurrent findings, from either RNA or DNA-based analyses, that may lead to discovery of actionable tumor dependencies. Such approaches, however, do not work on individual samples, since statistical significance cannot be assessed with an N of 1. Single cell analyses can potentially bypass these limitations by providing large tumor cell “cohorts” that can be studied using the same methodological approaches used for large cohorts, with the notable advantage that individual subpopulations—both transformed and stromal related—can be effectively identified, even from an individual sample. Unfortunately, by making ≥80% of the genes undetectable, single cell RNA-seq profiles present a different set of challenges. Indeed, most studies to date have been effective in stratifying tumors into some of the individual subpopulations they comprise, yet far less successful in identifying novel mechanisms and dependencies for targeted therapy.

By providing accurate protein activity quantitation VIPER allows direct extension of bulk-tissue mRNA analyses to single cells, with minimal to no loss of accuracy and sensitivity. Indeed, we have shown that VIPER-based protein activity measurements from single cell scRNA-seq profiles compare favorably with antibody-based measurements, leading to discovery of biologically and clinically relevant subpopulations and mechanisms that would have otherwise eluded scRNA-seq or antibody panel analyses^15, 17–19^. This includes using clinically-relevant methodologies, such as OncoTarget and OncoTreat, for the prioritization of small molecule inhibitors targeting Master Regulator dependencies of transformed subpopulations, as identified by single cell analyses, including coexisting subpopulations with potentially orthogonal drug sensitivity.

Comprehensive VIPER-based single-cell analysis of a single tumor biopsy from a cholangiocarcinoma patient, effectively predicted complementary drug sensitivity of three distinct transformed subpopulations and clarified the immunosuppressive nature of the cholangiocarcinoma TME in a specific patient—including a remarkably high (10-fold) stromal to tumor cell ratio—with significant therapeutic implications for other patients.

Cholangiocarcinoma has been previously described as having a significant stromal component, particularly with respect to fibroblasts, resulting in a dense extracellular matrix^4^. Our analysis confirmed significant fibroblast infiltrate yet also identified distinct T-cell subpopulations comprising nearly half of the tumor mass. While such an extent of lymphocytic infiltration suggests immune checkpoint inhibitors as a potentially valuable adjunct to traditional standard-of-care treatments, this is challenged by the presence of multiple immunosuppressive subpopulations. Specifically, single cell analysis identified sizeable presence of mast cells, myeloid cells, B-cells, and a cluster of neutrophils actively expressing IL8. In studies of melanoma^33^ and prostate cancer^34^, the latter have been reported to correlate with poor clinical outcome via myeloid-derived suppressor cell accumulation, for which several inhibitors are currently in clinical trials^35^. Targeting the immunosuppressive subpopulations identified by our analysis may thus could provide effective mechanisms to synergize with immune checkpoint inhibitors in cholangiocarcinoma.

As discussed, the extremely low tumor cell fraction—accounting only for <10% of the total cell count—renders bulk-tissue analyses of these tumors both largely irrelevant and potentially misleading. Indeed, validated single cell-based predictions were not recapitulated in bulk-tissue. Indeed, drug predictions emerging from bulk-tissue analysis are more likely to target T cells and fibroblasts in the TME than transformed cells. In contrast, quantitative protein activity assessment helped identify three molecularly distinct, coexisting transformed subpopulations. These were characterized by enrichment of KRAS signaling, TNFα signaling via NF-κB and EMT pathway, and MYC signaling hallmarks, respectively. Consistently, while clinically-relevant MR-inverter drugs were predicted for each subpopulation, only a handful were predicted to elicit sensitivity across multiple subtypes, of which two—dacinostat and plicamycin— were validated *in vivo*.

This level of tumor cell heterogeneity is previously unreported in cholangiocarcinoma studies. Yet, it represents an important characteristic of the disease that may explain failure of conventional treatment due to selection of drug-resistant subpopulations and suggests that effective treatment of this specific cholangiocarcinoma may require targeting all three subpopulations. Consistent with this perspective, plicamycin—predicted to target only two of the three, as confirmed *in vivo*—exhibited markedly reduced tumor growth control, compared to dacinostat, which targeted all three, albeit with differential sensitivity. Thus, our study provides potential rationales for the identification of drugs that may synergize by targeting independent transformed subpopulations.

While the extent to which the three transformed subpopulations identified by our analysis may generalize across other cholangiocarcinoma patients remains to be determined, we and others have shown that, at the single cell level, stable tumor cell states are much more conserved across patients than previously suspected^36, 37^. This is especially obvious when considering the activity of Master Regulators responsible for homeostatically controlling the transcriptional state of tumor cell, rather than gene expression. The latter can introduce significant confounding effects due to the presence of large, patient-specific copy number alterations that bias gene expression profiles while producing virtually identical Master Regulator activity signatures^24^.

The ability to create pseudo-bulk profiles, once the individual subpopulations have been identified by protein activity-based cluster analysis, further improves the sensitivity, accuracy, and reproducibility of drug predictions. Indeed, drug sensitivity and drug target predictions at the single-cell level were recapitulated and improved based on the analysis of synthetic bulks from each of the three transformed cell clusters. Furthermore, the PDX model generated from the same patient confirmed the differential activity of proteins in the three clusters, even though it could not differentiate between them. Of the four predicted drugs, dacinostat and plicamycin significantly decreased tumor growth rate vs. vehicle control and improve disease control rate *in vivo*. Plicamycin, predicted to strongly inhibit the two largest clusters but not the smallest, MYC activated one, slowed down tumor growth rate but did not have sustained effects, with only one out of eight mice showing disease control at 28 days. In contrast, dacinostat—predicted and validated to target all three cell clusters, albeit with variable sensitivity—demonstrated stable tumor size with no significant growth from baseline, at 28 days, for the majority of treated mice (4 of 6 mice). Given the paucity of available treatments for this disease, this provides a potentially relevant candidate for follow-up clinical studies.

In a prior study by Li et. al. of in vitro manual drug screening on 27 organoids derived from 3 cholangiocarcinoma and 2 hepatocellular carcinoma patients, plicamycin was found to be pan- effective across a broad range of organoids^38^. Out of 129 drugs screened in vitro, however, the authors found only 9 (7%)—across 5 classes of antineoplastic agents, including HDAC, proteasome, DNA topoisomerase II, protein translation, and RNA synthesis inhibitors— producing ≥ 90% killing across all organoids. Yet none of these were validated *in vivo* or demonstrated cholangiocarcinoma-specific activity at the tested concentrations.

Consistent with our unbiased, mechanism-based findings, plicamycin was one of the drugs that reduced organoid viability *in vitro*, suggesting potential value across a range of cholangiocarcinoma patients, with implications for future clinical trials. In contrast, however, two of four drugs predicted by our first-principles approach (50%) were validated *in vivo*. These were selected from a potential repertoire of >300 drugs whose mechanism of action had been dissected in the PanACEA database, thus representing a much-improved success rate compared to organoid-based drug screening, especially considering that none of the drugs in the prior study were validated *in vivo*. We use the term “mechanism-based” because we assess aberrant activity of candidate targets based on their ability to physically control the transcriptional state of the tumor cells, via their regulons; consistently, their MR-inversion potential is assessed from their mechanism of action, as also assessed by regulon-based assessment of the differential protein activity in drug vs. DMSO-treated cells.

Dacinostat, which we predicted and validated as even more effective than plicamycin, was not included in the drug panel assessed by Li et al^38^, thus representing a *bona fide* novel candidate for cholangiocarcinoma therapy. Consistently, CG200745, another HDAC inhibitor, was previously found to induce anti-tumor effects in cholangiocarcinoma cell lines via miRNAs targeting the Hippo pathway^39^. HDACs are also known to have a role in cholangiocarcinoma carcinogenesis and are being actively explored as therapeutic candidates, but each HDAC inhibitor has highly variable binding affinities for the different HDAC proteins^40^, with potential for very different therapeutic efficacy based on broad downstream transcriptional effects, which we directly assess by OncoTreat. To our knowledge, this is the first report of dacinostat as an HDAC inhibitor effective in cholangiocarcinoma treatment, *in vivo*. While these findings are limited to the patient assessed in the case study due to difficulty of cholangiocarcinoma PDX- engraftment, but may generalize across patients, similarly to plicamycin, meriting further clinical follow-up alone or in combination with current treatments, particularly given the dismal treatment outcomes and lack of response to current standard-of-care therapy.

In terms of limitations, it should be noted that drug sensitivity predictions are still not 100% accurate. Indeed, only 2 out of 4 drugs validated *in vivo*. This is the result of many potential causes, including (a) drug pharmacokinetics that prevent it from achieving the effective concentration in the tumor. This may be the case for flavopiridol, for instance, which has been reported to have adequate kinetics only in hematopoietic malignancies and not in solid tumors (personal communication), (b) mechanisms of cell adaptation that buffer the initial drug mechanism of action, as assessed at 24h from PanACEA, (c) false positive predictions common to all machine learning based algorithms. Still, the ability to reduce >300 drugs to a handful of candidates with 50% effectiveness *in vivo*, especially when considering the predicted subpopulation specific effects of each drug, is so far unmatched in the literature. Moreover, single cell analyses may fail to identify very rare, molecularly distinct, drug-resistant subpopulations with tumor initiating potential. This may require analyzing minimal residual tumor mass, enriching for the presence of these cells or flow sorting-based enriching using appropriate molecular markers, when available.

Finally, we have shown that Master Regulator proteins control not only the state of the tumor cell but also to implement a highly immunosuppressive TME^23^. As a result, the opportunity to use dacinostat or plicamycin in combination with immune checkpoint inhibitors is intriguing. In the future, we plan to extend these single cell-based approaches to a larger cohort of cholangiocarcinoma patients to identify drugs that may emerge as significant across multiple patients. In addition, our results suggest that these methodologies may be relevant to identify potential treatment for patients with rare, aggressive tumors that have not yet benefited from the large-cohort-based approaches used for more common malignancies.

## Methods

### Human Research Participation and Clinical Course of Treatment

Fresh, chemotherapy naïve surgical tissue was obtained with patient’s consent from central hepatectomy performed at tumor stage pT3N1M1. Tissue was dissociated immediately for scRNA-seq profiling. Following data analysis, tumor composition and OncoTreat/OncoTarget drug predictions were communicated to the medical team. Treatment with combination gemcitabine, cisplatin, and paclitaxel was initiated on a 21-day schedule one month after surgery, with dosing at day 1 and day 8, and CA19-9 tumor marker levels were drawn to monitor progression. With the patient’s consent, a trial of nivolumab was initiated following non-response to triple-combination chemotherapy and discovery of significant tumor T-cell infiltration by single-cell RNA Sequencing. Following interval improvement in CA19-9, the patient was enrolled in an ongoing trial of TP-1287, a formulation of flavopiridol with improved solid tumor PK (NCT03604783), but was able to receive only the first dose. Research was conducted in accordance with the Declaration of Helsinki.

### Tissue Dissociation

Fresh tumor tissue was minced to 2-4 mm sized pieces in a 6-cm dish and subsequently digested to single cell suspension using the Multi Tissue Human Tumor Dissociation Kit 1 (Miltenyi Biotec) and a gentleMACS OctoDissociator (Miltenyi Biotec) according to the manufacturer’s instructions. Dissociated cells were aliquoted for single-cell sequencing.

### Single-Cell RNA-Sequencing

Dissociated cells were processed for single-cell gene expression capture (scRNA-seq) using the 10X Chromium 3’ Library and Gel Bead Kit (10x Genomics), following the manufacturer’s user guide at the Columbia University Genome Center. After GelBead in-Emulsion reverse transcription (GEM-RT) reaction, 12-15 cycles of polymerase chain reaction (PCR) amplification were performed to obtain cDNAs used for RNA-seq library generation. Libraries were prepared following the manufacturer’s user guide and sequenced on Illumina NovaSeq 6000 Sequencing System. Single-cell RNA-seq data were processed with Cell Ranger software at the Columbia University Single Cell Analysis Core. Illumina base call files were converted to FASTQ files with the command “cellranger mkfastq.” Expression data were processed with “cellranger count” on a pre-built human reference set of 30,727 genes. Cell Ranger performed default filtering for quality control, and produced a barcodes.tsv, genes.tsv, and matrix.mts file containing transcript counts for each cell, such that expression of each gene is in terms of the number of unique molecular identifiers (UMIs) tagged to cDNA molecules corresponding to that gene. These data were loaded into the R version 3.6.1 programming environment, where the publicly available Seurat package was used to further quality-control filter cells to those with fewer than 25% mitochondrial RNA content, more than 1,000 unique UMI counts, and fewer than 25,000 unique UMI counts. Pooled distribution of UMI counts, unique gene counts, and percentage of mitochondrial DNA after QC-filtering is shown in Figure 2A.

### Single-cell RNA-seq Gene Expression Analysis

The Gene Expression UMI count matrix was processed in R using the Seurat SCTransform command to perform a regularized negative binomial regression, based on the 3,000 most variable genes. ScRNA-seq profiles were then clustered by a Resolution-Optimized Louvain Algorithm, leveraging mean Silhouette Score of individual profiles to select optimal resolution and avoid over-clustering^16^. Within each cluster MetaCells were computed for downstream regulatory network inference by summing SCTransform-corrected template counts for the 10 nearest neighbors of each cell, as assessed by Pearson correlation analysis. The resulting scRNA-seq dataset, representing 2,738 cells, was projected into its first 50 principal components using the RunPCA function in Seurat, and further reduced to a 2-dimensional visualization space using the RunUMAP function, with method umap-learn and Pearson correlation as the distance metric between cells. Differential Gene Expression between clusters was computed by the MAST hurdle model for single-cell gene expression, as implemented in the Seurat FindAllMarkers command, with log fold change threshold of 0.5 and minimum fractional expression threshold of 0.25, indicating that the resulting gene markers for each cluster are restricted to those with log fold change greater than 0 and non-zero expression in at least 25% of the cells in the cluster.

### Semi-Supervised Cell Type Calling

Unbiased inference of cell types was performed using the SingleR package and the preloaded Blueprint-ENCODE reference, which includes normalized expression values for 259 bulk RNA-seq samples generated by Blueprint and ENCODE from 43 distinct cell types, representing pure, non-transformed cell populations^41^. The SingleR algorithm computes correlation between each individual cell and each of the 259 reference samples, and then assigns a label of the cell type with highest average correlation to the individual cell and a p-value computed by Wilcox’s test of correlation to that cell type compared to all other cell types. Unsupervised clusters, as determined by the resolution-optimized Louvain algorithm, were consensus-labelled according to the most highly represented SingleR cell type label within that cluster, among all labels with *p* < 0.05.

### Copy Number Inference

Copy Number Variation (CNV) was inferred from gene expression counts at the single cell level using the InferCNV package. Cells were clustered according to their unsupervised clustering label by protein activity, and the large T-cell cluster was used as a Copy-Number-Normal reference set. The 140-cell epithelial cluster, as labelled by SingleR, was confirmed to exhibit significant copy-number alteration across the entire genome relative to other cell types, confirming its transformed nature.

### Regulatory Network Inference

Within each gene expression cluster, MetaCells were computed by summing SCTransform-corrected template counts for the 10 nearest neighbors of each cell by Pearson correlation distance. For clusters exceeding 200 cells, MetaCells were randomly sub-sampled to *n* = 200. For each cluster a transcriptional regulatory network was inferred by ARACNe algorithm analysis^42^. ARACNe was run with 200 bootstrap iterations using 1,785 transcription factors—including genes annotated as GO:0003700 (transcription factor activity), GO:0003677 (DNA binding), GO:0030528 (transcription regulator activity), GO:0003677 (DNA-interacting), or GO:0045449 (regulation of transcription) in the Gene Ontology database^43^—668 transcriptional cofactors—a manually curated list, not overlapping with the transcription factor list, based on genes annotated as GO:0003712 (transcription cofactor activity), GO:0030528 (transcription regulator activity), or GO:0045449 (regulation of transcription, DNA-templated)—3,455 signaling pathway related genes—annotated as GO:0007165 (signal transduction), and either GO:0005622 (intracellular) or GO:0005886 (plasma membrane)—and 3,620 surface markers—annotated as GO:0005886 or GO:0009986 (cell surface). Parameters were set as follows: DPI = 0 (Data Processing Inequality) tolerance and MI (Mutual Information) p-value threshold p ≤ 10^-8^, computed by shuffling the original dataset as a null model. Each gene list used to run ARACNe is available on Github, along with the generated ARACNe tables.

### Protein Ahctivity Inference

Tissue specific regulons were inferred directly from the analysis of each single-cell subpopulations, as inferred by gene expression-based clustering, using ARACNe^42^, an information theoretic algorithm that has been experimentally validated in dozens of tissue contexts, with >70% target identification accuracy^44^. This is because lineage related cells have highly conserved regulatory networks^45^, such that even rough clustering, as produced by gene expression, is sufficient. By integrating the expression of the most statistically significant 100 transcriptional targets in each regulon, VIPER can effectively measure protein transcriptional activity, including for protein whose encoding gene is undetected in scRNA-seq data, thus virtually eliminating gene dropouts^46^. A fixed number of gene is selected to avoid set size-related bias in GSEA.

Protein activity was first inferred for all cells by running the metaVIPER algorithm with all subpopulation-specific ARACNe networks on the SCTransform-scaled single-cell gene expression signature. Because the SCTransform-scaled gene expression signature is already internally normalized to maximize differences among individual subpopulations, VIPER normalization option was set to “none.” The resulting VIPER matrix included 1,602 proteins whose differential activity, compared to the average of all cells, was successfully inferred for all 2,738 cells. Focusing only on transformed cells, this protein activity sub-matrix was loaded into a Seurat Object with CreateSeuratObject, then projected into its first 50 principal components using the RunPCA function in Seurat, and further reduced into a 2-dimensional visualization space using the RunUMAP function with method umap-learn and Pearson correlation as the distance metric between cells. Differential Protein Activity between clusters, as identified by resolution-optimized Louvain, was computed using a bootstrapped t-test, with 100 bootstraps. Top proteins for each cluster were ranked by p-value (Figure 4C). Activity-based analysis identified three molecularly distinct clusters of transformed cell, which could not be stratified by gene expression-based clustering.

### Single-cell OncoTarget Analysis

To identify aberrantly activated, actionable proteins in each of the three transformed cell subpopulations, we generated single cell-normalized differential gene expression signatures by comparing the log-Normalized Counts-Per-Million (CPM) of each gene in each single cell (log10[CPM-UMI count +1]) to the log-Normalized Transcripts-Per-Million (TPM) data (log10[TPM+1]) of the entire publicly available TCGA repository, as an effective universal reference to assess the differential expression of regulon genes. VIPER analysis of these gene expression signatures—using each cell’s tissue-matched, single-cell ARACNe network—was used to produce a single cell-specific differential protein activity matrix. For OncoTarget analysis, only proteins identified as established high-affinity binders (≤ 1μM) of FDA-approved or late-stage investigational drug compounds in DrugBank were considered. The Normalized Enrichment Score (NES) of each protein, in each cell, as assessed by GSEA of its regulon genes in differentially expressed genes, using the aREA algorithm^30^, was then converted to a Bonferroni-corrected p-value. Proteins with median *p* ≤ 10^-5^ in any tumor cell sub- cluster were selected as candidate actionable targets.

To assess robustness of OncoTarget predictions, the same analysis was performed on a synthetic bulk profile generated for each transformed cell sub-cluster, thus trading single cell resolution for RNA-seq profile depth, by summing UMI counts across all cells in the same cluster. Confirming our VIPER reproducibility-based expectations, this produced the exact same druggable protein predictions as the integration of predictions from each individual cell in a cluster (Figure 5D).

### Single-cell OncoTreat Analysis

To improve drug predictions by assessing the proteome-wide mechanism of action of candidate drugs, including off-target and downstream effector proteins, we leveraged PanACEA^31^, a database comprising the RNA-seq profiles of 16 cell lines— including BT20, HCC1143, HSTS, KRJ1, IOMM, U87, HF2597, H1793, ASPC1, PANC1, LNCAP, DU145, TCCSUP, EFO21, ASPC1, and PANC1, at 24h following perturbation with >300 drugs and vehicle control (DMSO). As for OncoTarget, we computed single-cell protein activity based on their differential gene expression compared to the entire TCGA repository.

Regulatory networks fidelity to cholangiocarcinoma was assessed based on their ability to identify statistically significantly differentially active proteins (Master Regulators). As shown in^47^, the rationale is that suboptimal networks can only decrease but not increase the statistical significance of differentially active proteins. To assess fidelity, Normalized Enrichment Scores representing differential activity of each protein *p_i_* in each single cell *c_j_*, using each available network model *N_k_*, were computed and converted to Bonferroni-corrected p-values. Then for each single cell *c_j_* and network model *N_k_*, the -log_10_ *p* of all significantly differentially active proteins (*p* ≤ 10^-2^) were added:

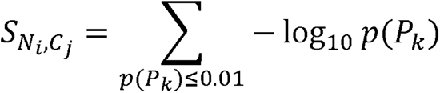

Finally, the network model-specific median M(*N_i_*) of the resulting distribution *p*(*s_Ni_,C_j_*)*j*=1,…,*n*, across all *n* single cells, was computed. The pancreatic adenocarcinoma (PDA) network was selected as the one with the highest median score among all available tumor-specific network models (Supplementary Figure 1A), consistent with the expected similarity between transformed bile duct and pancreatic duct epithelial cells. Based on this result, we assessed the fidelity of the two PDA cell lines in PanACEA—ASPC1 and PANC1, for which drug perturbation profiles were available—and the transformed single cells from cholangiocarcinoma sample, by assessing the enrichment of the top 50 and bottom 50 most differentially active proteins (MRs) in each transformed single-cell in protein differentially active signature in each cell line (OncoMatch method^48^), integrated across all single cells using Fisher’s method (Supplementary Figure 1B). The analysis identified ASPC1 as a very high-fidelity model (*p* < 10^-15^) significantly outperforming PANC1 (*p* = 0.99), resulting in its selection for follow-up OncoTreat analysis.

ASPC1-based OncoTreat analysis was then used to assess the statistical significance of each drugs in terms of inverting the activity of Master Regulator proteins inferred either from individual scRNA-seq or synthetic bulk RNA-seq profiles, based on the enrichment of Master Regulator proteins in proteins differentially active in drug vs. DMSO-treated ASPC1 cells, with negative NES values corresponding to MR activity inversion. Statistical significance as assessed from individual cells was integrated using Fisher’s method, such that drugs predicted to invert the MR activity of a larger number of single cells were more highly ranked. In Figure 5B we show the results of these analyses, including the integrated p-value across all tumor cells for the 25 most statistically significant MR-inverter drugs, the 25 most significant drugs based on OncoTreat analysis of the synthetic bulk from each cluster, and the 25 most significant drugs for each individual cluster. As for OncoTarget, synthetic bulks profiles were generated by summing the UMI counts of each gene across all same-cluster cells.

### OncoTarget/OncoTreat Analysis of PDX Bulk RNA-Seq Data

A Bulk RNA-Seq profile was generated from tumor samples harvested from the pre-treatment PDX model and analyzed using the same OncoTarget and OncoTreat methodologies used for the synthetic bulk samples, including use of the same PDAC regulatory model and ASPC1 cell line. As described previously, a differential gene expression signature was computed by scaling the log-transformed TPM data to the average of TCGA and then analyzed by VIPER to generate a protein activity matrix. Enrichment of Master Regulators identified from analysis of the three transformed subpopulations in proteins differentially active in the PDX model was assessed by the aREA implementation of GSEA. Figure 6A shows that the PDX analysis recapitulated several subpopulation-specific MRs, as identified by single cell analysis, with higher enrichment in MRs from the two largest subpopulations, associated with KRAS and Nf-kb/EMT signaling, respectively. However, as expected, MR specificity was lost, such that which drug would likely elicit sensitivity in which subpopulation may not be determined.

### PDX Model Establishment and Treatment

Mice were maintained under barrier conditions and experiments were conducted using protocols and conditions approved by the Memorial Sloan Kettering Cancer Center (MSKCC) Institutional Animal Care and Use Committee (IACUC) under protocol 16-08-011. Patient derived tumor tissue to generate PDX models were obtained under the MSKCC Institutional Review Board (IRB)-approved protocols #17-387 and #06-107. PDX mouse models were established by implanting tumor cells subcutaneously into non-obese diabetic/severe combined immunodeficiency interleukin-2R gamma null, HPRT null (NSGH) mice (Jackson Labs, IMSR Cat# JAX:012480, RRID: IMSR_JAX:012480). Mice were assigned into five treatment groups using block randomization and treatment initiated when tumor volume (TV) was ∼100 mm^3^. Tumor size was assessed by caliper measurement, twice weekly, and TV was calculated as follows: TV = width^2^ X ½ length. Mice were assigned to the following treatment arms (n=8/arm): (1) vehicle control, (2) plicamycin 0.2 mg/kg IP 3 times/week, (3) AT9283 15 mg/kg daily, IP 4 days on 3 days off, (4) flavopiridol 15 mg/kg PO daily, 5 days on 2 days off, and (5) dacinostat 25 mg/kg IP daily. All animals were treated for at least 4 weeks with tumor measurements continued beyond treatment to determine time to treatment failure. Prior to treatment efficacy studies, drug dose and schedules were evaluated in tumor-naïve animals to confirm tolerability of each drug for the duration of the treatment period. Treatment failure was defined as >100% increase in tumor volume relative to baseline in each respective mouse (i.e., tumor doubling). For *in vivo* statistical analysis, the Mann-Whitney-Wilcoxon method was used to evaluate differences in distribution of tumor volume between treatment groups. Vardi’s test was used to evaluate differences in the area under the curve (AUC) between treatment groups. Disease control rate was defined as the percentage of mice that did not satisfy criteria for treatment failure (progressive disease) for the duration of the study period (observation up to Day 45). Kaplan-Meier survival curves were compared using the log-rank test. Statistical analysis was performed using R software (v3.5.0). Waterfall plots and tumor volume curves for in vivo analysis were generated with GraphPad Prism (v8.4.1). Statistical significance was defined as *p* ≤ 0.05.

### Single-Cell RNA-Sequencing of Post-Treatment PDX Models

For the dacinostat, plicamycin, and vehicle control—the first two achieving statistically significant tumor growth control—fresh tumor tissue was collected at the study endpoint from one animal and dissociated to generate scRNA-seq profiles. The same protocol and analysis methodology described above was used, i.e. the combination of ARACNe and VIPER algorithms.

The transformed, human-derived nature of profiled cells was confirmed by InferCNV and by consensus alignment of RNA reads to the Human vs. mouse reference genome (Figure S2). Once isolated (Figure 7A), the similarity of post-treatment tumor cells from each mouse to each patient-derived subpopulation (Clusters *C*_1_, *C*_2_, and *C*_3_) was assessed by aREA-based GSEA. This yielded matched post-treatment PDX tumor cell population for each of the three clusters, as well as a small newly observed population with significant enrichment for both cluster *C*_1_ and *C*_2_, which may correspond to doublets (Figure 7A). Fractional abundance of each post- vs. pre-treatment transformed cell population was assessed in each animal, and log-fold-change in frequency of each population in dacinostat-treated and plicamycin-treated mice was assessed relative to vehicle control, normalized to overall tumor growth reduction, to determine whether OncoTreat correctly predicted subpopulation-specific drug sensitivity (Figure 7B-C).

## Acknowledgements

This work was supported by the NCI Cancer Target Discovery and Development Program (U01 CA217858), an NCI Outstanding Investigator Award (R35 CA197745), and NIH Shared Instrumentation Grants (S10 OD012351 and S1 0OD021764), all to Andrea Califano. Also, this research was funded in part through the NCI Cancer Center Support Grant (P30 CA013696).

## Author Contributions

A.O. and L.T. conceived of and implemented the computational analyses, and selected candidate drugs for follow-up validation, F.D.C. and D.Y. coordinated PDX model engraftment and treatment with candidate drug compounds. E.F., S.B., and Y.S. recruited the patient and coordinated tissue acquisition for single-cell RNA-Sequencing. A.O. coordinated and performed single-cell RNA-Sequencing of on-treatment PDX samples. A.C. conceived of the core algorithms, and guided computational analyses and validation design.

---

**Supplementary Figure 1:**
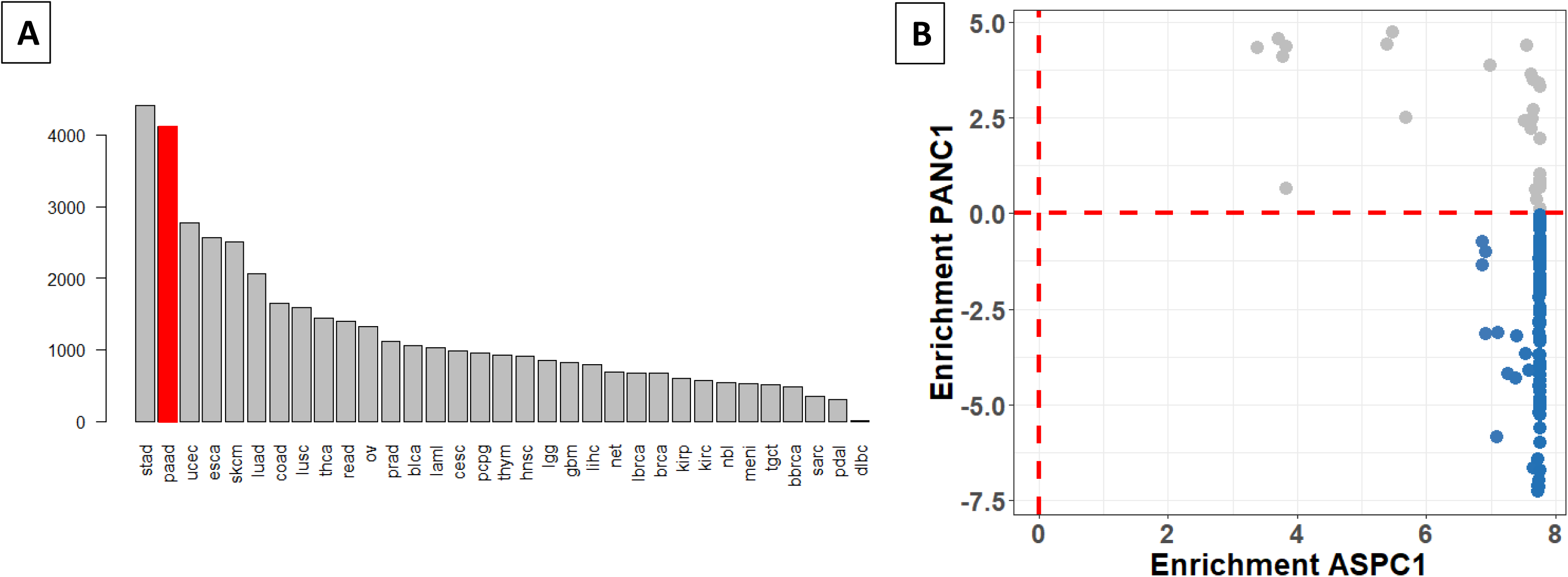
**(A)** Results of the network-fidelity assessment to determine the best ARACNe-inferred regulatory model for the OncoTreat analysis of tumor cells. The top-matched network for which we have perturbational data is the Pancreatic Adenocarcinoma Interactome (second one in the overall ranking shown in red) **(B)** GSEA of the top 50 and bottom 50 most differentially active proteins in each single-cells (Master Regulators) in proteins differentially active in ASPC1 and PANC1 cells shows that virtually all tumor cells were enriched in the ASPC1 signature while most were negatively enriched in the PANC1 signature, leading to ASPC1 selection for OncoTreat analysis.

**Supplementary Figure 2:**
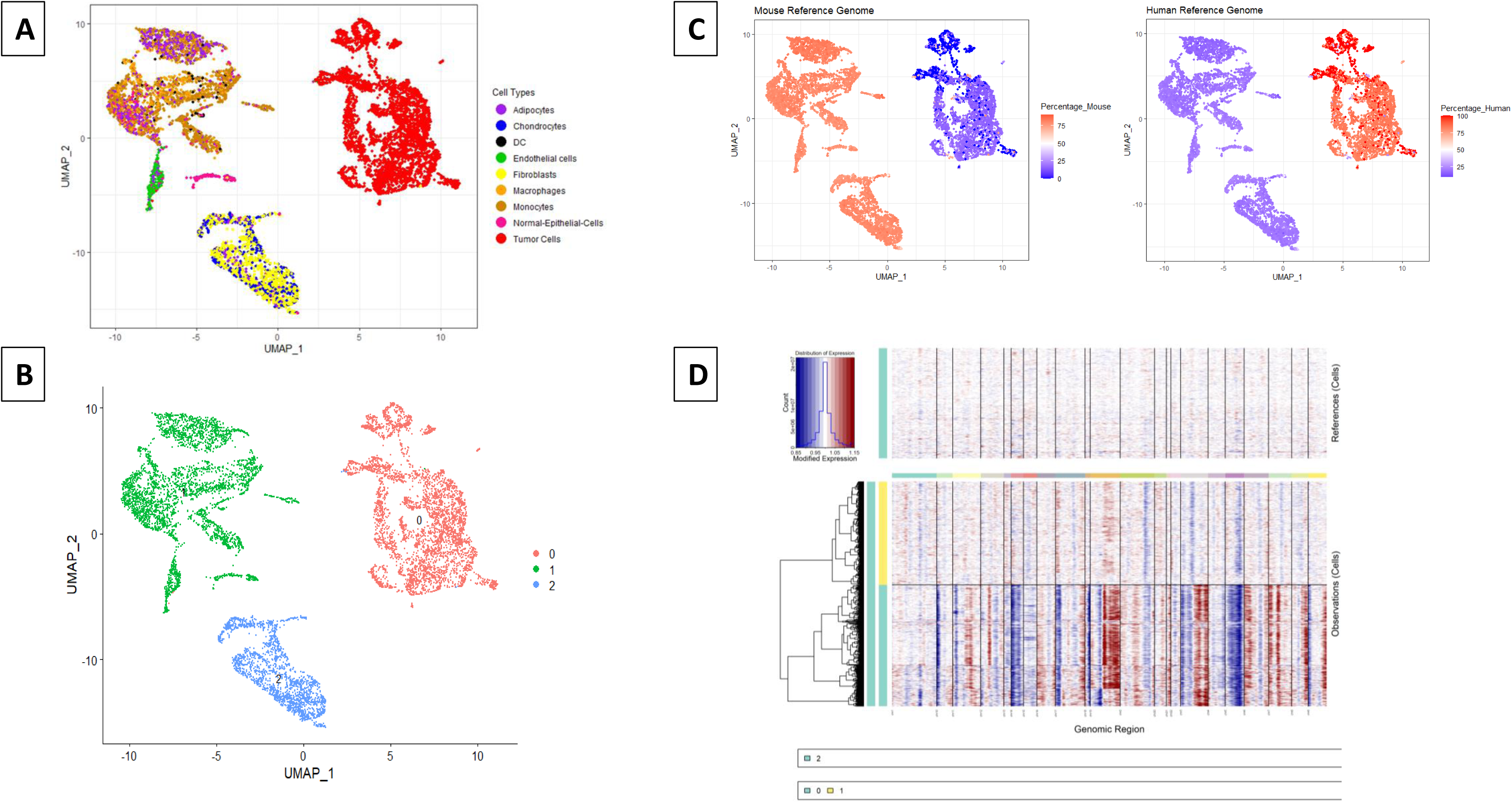
**(A**) Protein activity-based UMAP projection showing combined single-cell RNA Sequencing data in post-treatment PDX mice treated with dacinostat, plicamycin and vehicle control. Cell type identity was inferred by SingleR analysis **(B)** Protein activity-based unsupervised clustering analysis. **(C)** Heat plot showing proportion of reads from PDX model mapped to the mouse genome reference (left) and to the human genome reference (right). **(D)** InferCNV showing that extensive copy number alterations are limited to cluster 0

**Supplementary Figure 3:**
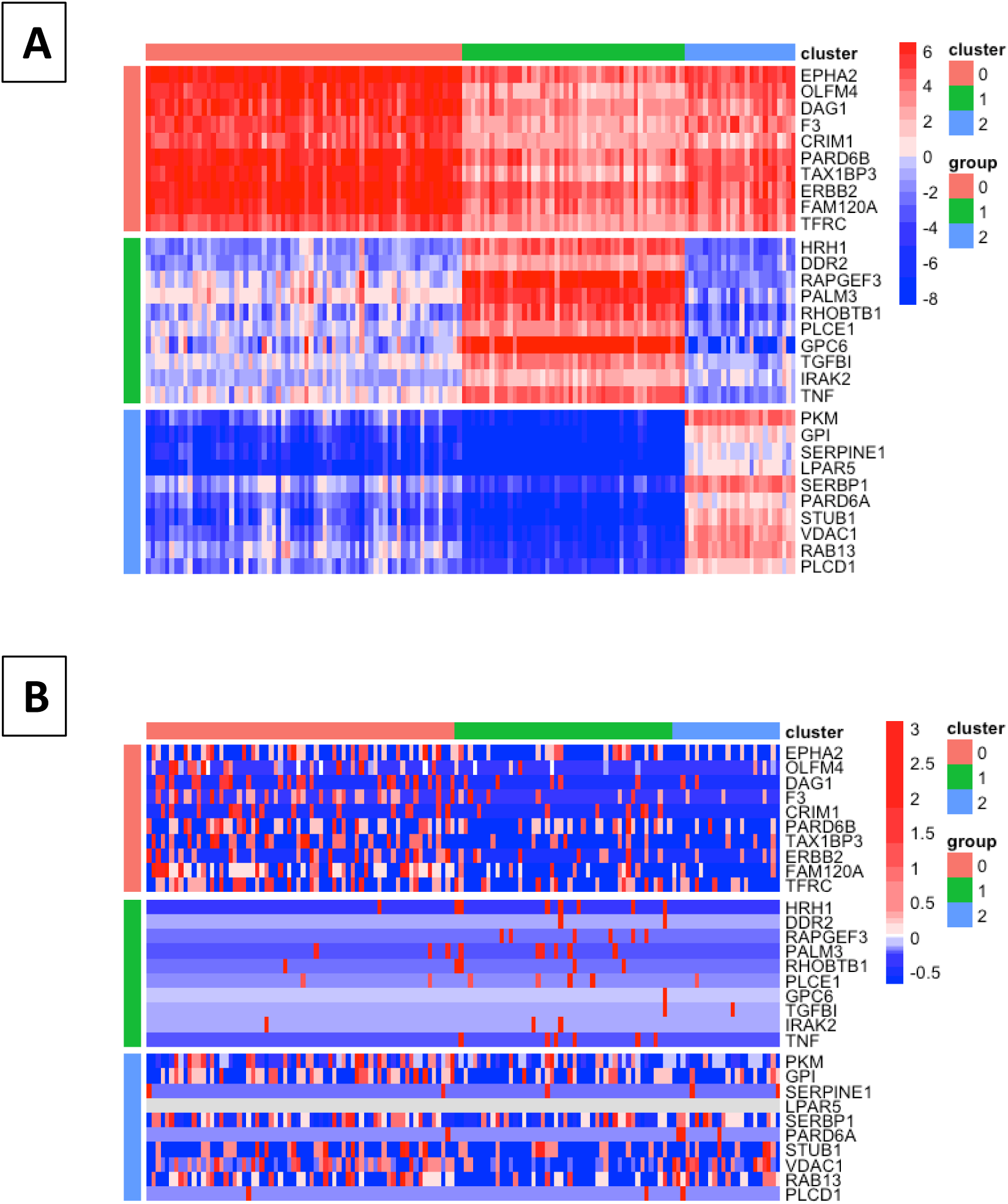
**(A)** Heatmap of the 10 most differentially active surface marker proteins distinguishing the three transformed cell subpopulations, by VIPER analysis. **(B)** Gene expression heatmap for the gene encoding for proteins shown in S3A shows dramatic gene dropout effects.

**Supplementary Figure 4:**
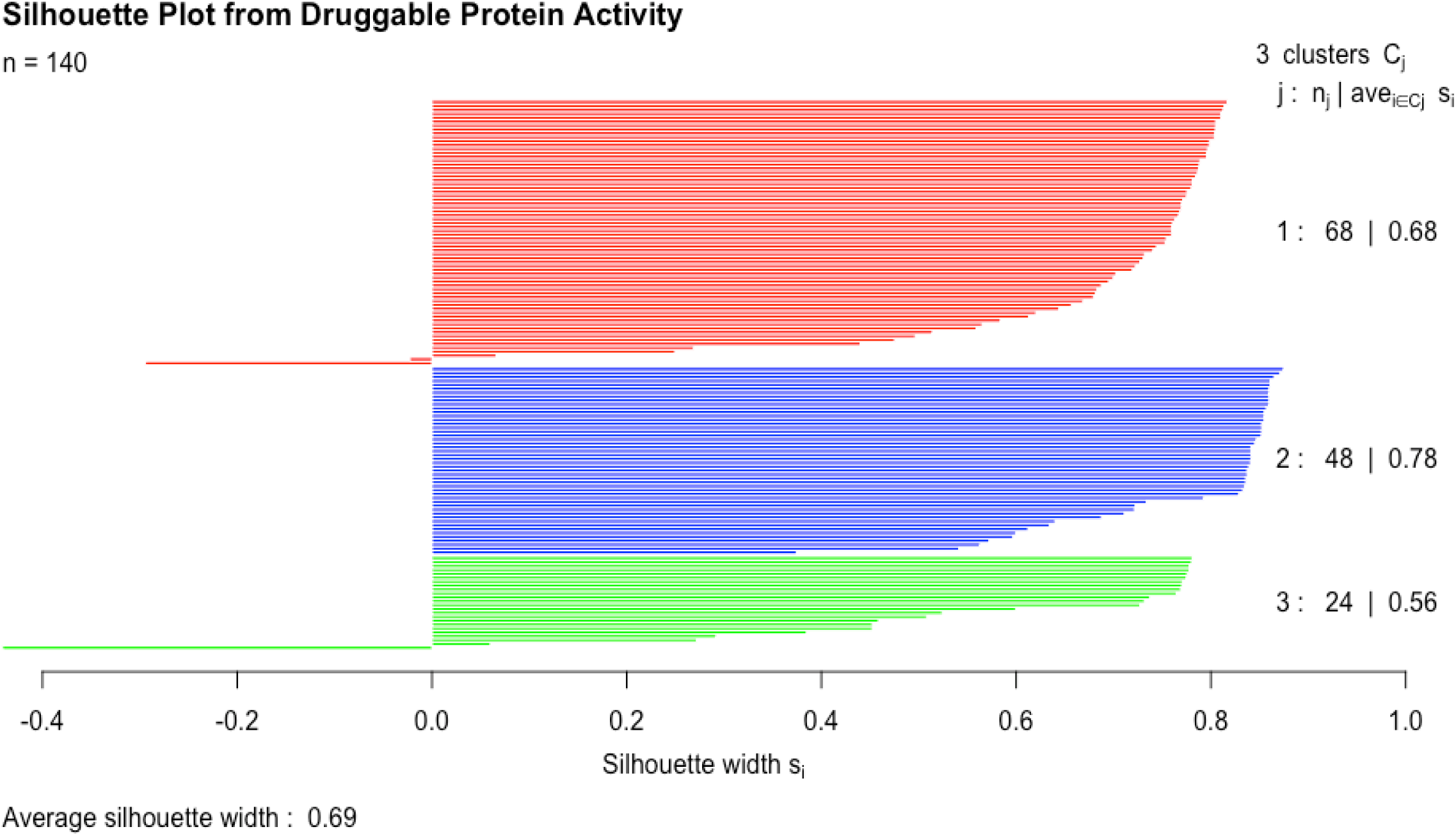
Silhouette score analysis shows that individual transformed cells are confidently assigned to their respective clusters, with distance computed by Pearson correlation on VIPER activity of the top statistically significant druggable proteins (average *p* ≤ 10^-5^ across each cluster).

**Supplementary Figure 5:**
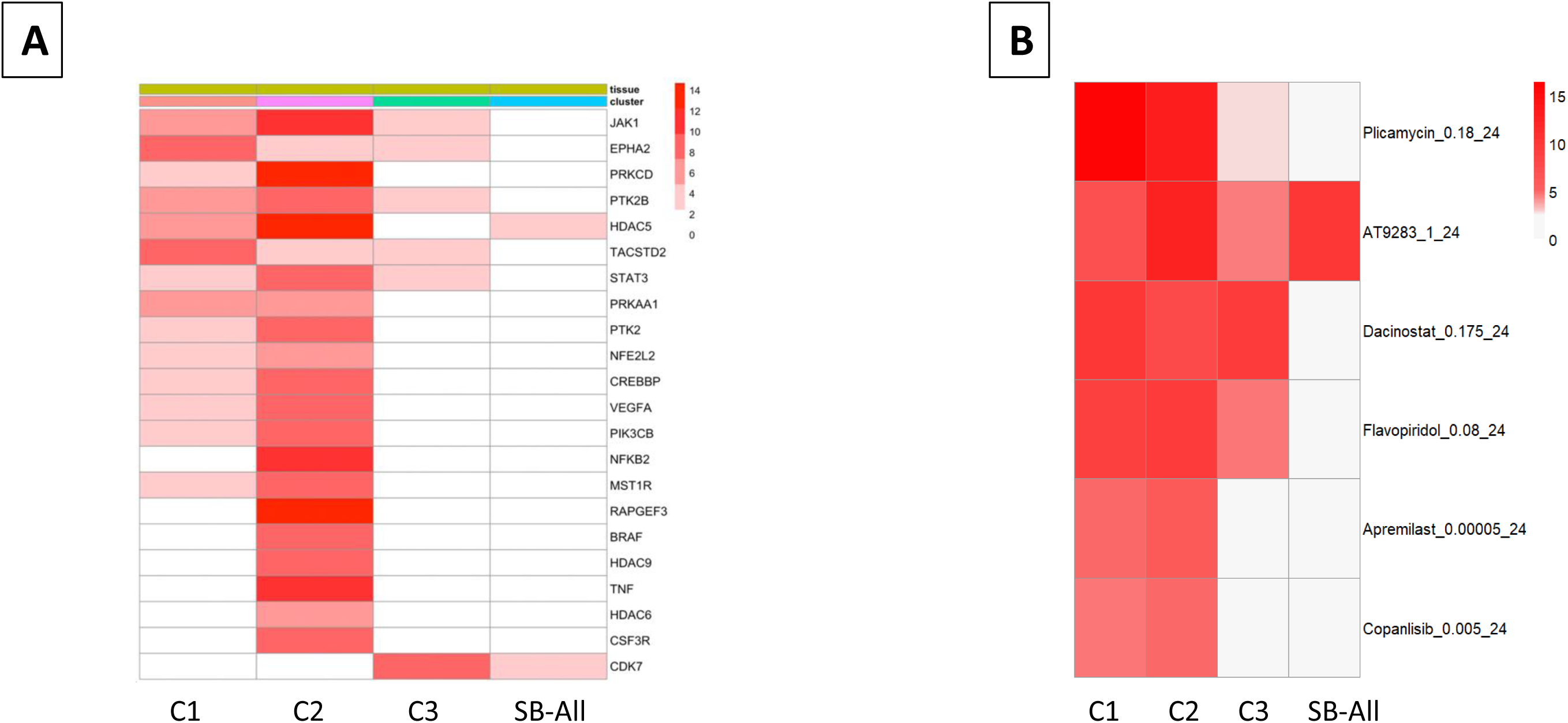
**(A)** Most significant activated actionable proteins as predicted by OncoTarget from synthetic bulk samples generated from each transformed cell subpopulation and from the synthetic bulk representing the entire sample, including all cells both tumor and stroma-related (right-most column). **(B)** Top predicted drugs by OncoTreat for the same set of synthetic bulk samples, with associated statistical significance (-log_10_(*p*)).

## References

1. Brunt E, Aishima S, Clavien PA, et al: cHCC-CCA: Consensus terminology for primary liver carcinomas with both hepatocytic and cholangiocytic differentation. Hepatology 68:113–126, 2018

2. Dhanasekaran R, Hemming AW, Zendejas I, et al: Treatment outcomes and prognostic factors of intrahepatic cholangiocarcinoma. Oncology reports 29:1259–1267, 2013

3. Society AC: Survival Rates for Bile Duct Cancer, 2021

4. Fabris L, Perugorria MJ, Mertens J, et al: The tumour microenvironment and immune milieu of cholangiocarcinoma. Liver International 39:63–78, 2019

5. Boscoe AN, Rolland C, Kelley RK: Frequency and prognostic significance of isocitrate dehydrogenase 1 mutations in cholangiocarcinoma: a systematic literature review. J Gastrointest Oncol 10:751–765, 2019

6. Finak G, McDavid A, Yajima M, et al: MAST: a flexible statistical framework for assessing transcriptional changes and characterizing heterogeneity in single-cell RNA sequencing data. Genome biology 16:1–13, 2015

7. Zheng GX, Terry JM, Belgrader P, et al: Massively parallel digital transcriptional profiling of single cells. Nature communications 8:1–12, 2017

8. Stuart T, Satija R: Integrative single-cell analysis. Nature Reviews Genetics 20:257–272, 2019

9. Sade-Feldman M, Yizhak K, Bjorgaard SL, et al: Defining T cell states associated with response to checkpoint immunotherapy in melanoma. Cell 175:998–1013. e20, 2018

10. Jerby-Arnon L, Shah P, Cuoco MS, et al: A cancer cell program promotes T cell exclusion and resistance to checkpoint blockade. Cell 175:984–997. e24, 2018

11. Chung W, Eum HH, Lee H-O, et al: Single-cell RNA-seq enables comprehensive tumour and immune cell profiling in primary breast cancer. Nature communications 8:1–12, 2017

12. Neftel C, Laffy J, Filbin MG, et al: An Integrative Model of Cellular States, Plasticity, and Genetics for Glioblastoma. Cell 178:835–849 e21, 2019

13. Alvarez MJ, Shen Y, Giorgi FM, et al: Functional characterization of somatic mutations in cancer using network-based inference of protein activity. Nature genetics 48:838–847, 2016

14. Piovan E, Yu J, Tosello V, et al: Direct reversal of glucocorticoid resistance by AKT inhibition in acute lymphoblastic leukemia. Cancer Cell 24:766–76, 2013

15. Obradovic A, Chowdhury N, Haake SM, et al: Single-cell protein activity analysis identifies recurrence-associated renal tumor macrophages. Cell 184:2988–3005 e16, 2021

16. Obradovic A, Chowdhury N, Haake SM, et al: Single-cell protein activity analysis identifies recurrence-associated renal tumor macrophages. Cell 184:2988–3005. e16, 2021

17. Arumugam K, Shin W, Schiavone V, et al: The Master Regulator Protein BAZ2B Can Reprogram Human Hematopoietic Lineage-Committed Progenitors into a Multipotent State. Cell Rep 33:108474, 2020

18. Son JB, Ding H, Farb TB, et al: Reversibility of beta-cell failure in type 2 diabetes through BACH2 inhibition. Journal of Clinical Investigation in press, 2021

19. Elyada E, Bolisetty M, Laise P, et al: Cross-Species Single-Cell Analysis of Pancreatic Ductal Adenocarcinoma Reveals Antigen-Presenting Cancer-Associated Fibroblasts. Cancer Discov 9:1102–1123, 2019

20. Mundi PS, Dela Cruz FS, Grunn A, et al: Pre-clinical validation of an RNA-based precision oncology platform for patient-therapy alignment in a diverse set of human malignancies resistant to standard treatments. bioRxiv 2021.10.03.462951, 2021

21. Weinstein IB: Cancer. Addiction to oncogenes--the Achilles heal of cancer. Science 297:63–4, 2002

22. Zeleke T, Pan Q, Chiuzan C, et al: Network-based assessment of HDAC6 activity is highly predictive of pre-clinical and clinical responses to the HDAC6 inhibitor ricolinostat. medRxiv, 2020

23. Alvarez MJ, Subramaniam PS, Tang LH, et al: A precision oncology approach to the pharmacological targeting of mechanistic dependencies in neuroendocrine tumors. Nat Genet 50:979–989, 2018

24. Paull EO, Aytes A, Jones SJ, et al: A modular master regulator landscape controls cancer transcriptional identity. Cell 184:334–351 e20, 2021

25. Aran D, Looney AP, Liu L, et al: Reference-based analysis of lung single-cell sequencing reveals a transitional profibrotic macrophage. Nat Immunol 20:163–172, 2019

26. Loeuillard E, Yang J, Buckarma E, et al: Targeting tumor-associated macrophages and granulocytic myeloid-derived suppressor cells augments PD-1 blockade in cholangiocarcinoma. The Journal of clinical investigation 130, 2020

27. Rizvi S, Gores GJ: Pathogenesis, diagnosis, and management of cholangiocarcinoma. Gastroenterology 145:1215–1229, 2013

28. Wang P, Lv L: miR-26a induced the suppression of tumor growth of cholangiocarcinoma via KRT19 approach. Oncotarget 7:81367, 2016

29. Chang K, Creighton CJ, Davis C, et al: The Cancer Genome Atlas Pan-Cancer analysis project. Nature Genetics 45:1113–1120, 2013

30. Alvarez MJ, Shen Y, Giorgi FM, et al: Functional characterization of somatic mutations in cancer using network-based inference of protein activity. Nat Genet 48:838–47, 2016

31. Douglass EF, Allaway RJ, Szalai B, et al: A Community Challenge for Pancancer Drug Mechanism of Action Inference from Perturbational Profile Data. Cell Med Reports (in press), 2020

32. Rajbhandari P, Lopez G, Capdevila C, et al: Cross-Cohort Analysis Identifies a TEAD4-MYCN Positive Feedback Loop as the Core Regulatory Element of High-Risk Neuroblastoma. Cancer Discov 8:582–599, 2018

33. Tobin RP, Jordan KR, Kapoor P, et al: IL-6 and IL-8 are linked with myeloid-derived suppressor cell accumulation and correlate with poor clinical outcomes in melanoma patients. Frontiers in oncology 9:1223, 2019

34. Lopez-Bujanda ZA, Haffner MC, Chaimowitz MG, et al: Castration-mediated IL- 8 promotes myeloid infiltration and prostate cancer progression. Nature Cancer:1–16, 2021

35. Gonzalez-Aparicio M, Alfaro C: Significance of the IL-8 pathway for immunotherapy. Human vaccines & immunotherapeutics 16:2312–2317, 2020

36. Laise P, Turunen M, Maurer HC, et al: Pancreatic Ductal Adenocarcinoma Comprises Coexisting Regulatory States with both Common and Distinct Dependencies. bioRxiv 2020.10.27.357269, 2021

37. Ding H, Burgenske DM, Zhao W, et al: Single-cell based elucidation of molecularly-distinct glioblastoma states and drug sensitivity. bioRxiv, 2019

38. Li L, Knutsdottir H, Hui K, et al: Human primary liver cancer organoids reveal intratumor and interpatient drug response heterogeneity. JCI insight 4, 2019

39. Jung DE, Park SB, Kim K, et al: CG200745, an HDAC inhibitor, induces anti-tumour effects in cholangiocarcinoma cell lines via miRNAs targeting the Hippo pathway. Scientific reports 7:1–13, 2017

40. Pant K, Peixoto E, Richard S, et al: Role of histone deacetylases in carcinogenesis: potential role in cholangiocarcinoma. Cells 9:780, 2020

41. Martens JH, Stunnenberg HG: BLUEPRINT: mapping human blood cell epigenomes. Haematologica 98:1487, 2013

42. Basso K, Margolin AA, Stolovitzky G, et al: Reverse engineering of regulatory networks in human B cells. Nat Genet 37:382–90, 2005

43. Ashburner M, Ball CA, Blake JA, et al: Gene ontology: tool for the unification of biology. The Gene Ontology Consortium. Nat Genet 25:25–9, 2000

44. Basso K, Margolin AA, Stolovitzky G, et al: Reverse engineering of regulatory networks in human B cells. Nature genetics 37:382–390, 2005

45. Mani KM, Lefebvre C, Wang K, et al: A systems biology approach to prediction of oncogenes and molecular perturbation targets in B-cell lymphomas. Mol Syst Biol 4:169, 2008

46. Ding H, Douglass EF, Sonabend AM, et al: Quantitative assessment of protein activity in orphan tissues and single cells using the metaVIPER algorithm. Nature communications 9:1–10, 2018

47. Aytes A, Mitrofanova A, Lefebvre C, et al: Cross-species regulatory network analysis identifies a synergistic interaction between FOXM1 and CENPF that drives prostate cancer malignancy. Cancer Cell 25:638–51, 2014

48. Alvarez MJ, Yan P, Alpaugh ML, et al: Reply to ’H-STS, L-STS and KRJ-I are not authentic GEPNET cell lines’. Nat Genet 51:1427–1428, 2019

49. Hanahan D, Weinberg RA: Hallmarks of cancer: the next generation. Cell 144:646–74, 2011

